# DM4ISL: high-fidelity and uncertainty-aware in silico labeling of fluorescence images via diffusion models

**DOI:** 10.64898/2026.11.10.765545

**Authors:** Oded Rotem, Orit Kliper-Gross, Assaf Zaritsky

## Abstract

In silico labeling predicts organelle-specific localizations from label-free images, potentially enabling longitudinal and multiplexed live-cell imaging. Here we present DM4ISL (Diffusion Model for In Silico Labeling), a high-fidelity and uncertainty-aware model that overcomes current limitations, such as blurry or structurally inaccurate predictions, by resolving the intrinsic entanglement of biological signal and noise that characterizes fluorescence imaging. DM4ISL leverages the iterative nature of diffusion through a step-wise self-ensemble averaging approach that marginalizes these stochastic fluctuations, preventing the overfitting to noise that typically degrades standard final-step predictions. Benchmarking across six organelles demonstrates that DM4ISL systematically outperforms state-of-the-art architectures in structural fidelity, as confirmed through both pixel-based and application-specific metrics. Furthermore, calculating the standard deviation across intermediate denoising steps generates localized uncertainty maps that automatically detect erroneous predictions at inference time. Providing high-fidelity predictions alongside built-in reliability metrics, DM4ISL offers a robust method for trustworthy, data-driven discovery in cell biology.

## Introduction

Fluorescence microscopy enables mechanistic investigation of the cell’s organization, dynamics and function ^1^ but suffers from several drawbacks that hamper long-term rapid live imaging of multiple subcellular structures while maintaining cells in their native state. These drawbacks include disruption of cellular functions via the fluorescence labeling, limitation in the number of distinguishable fluorescent labels ^2^, photobleaching and phototoxicity ^3,4^. In silico labeling is a computational technique that predicts fluorescence-labeled images of specific organelles from label-free transmitted light microscopy images by training deep learning models on matched pairs of label-free and fluorescent images ^5^. This enables label-free, multiplexed, and less phototoxic live-cell imaging, offering a powerful tool for studying cellular structures and dynamics without physical fluorescent labeling ^6–8^. However, despite holding great promise, inherent difficulties hinder the full potential of in silico labeling for driving new biological discoveries. First, current in silico labeling models, based on U-Net and GANs ^6,7,9^, even with an attention mechanism ^10^, generate blurry fluorescent images that smooth out fine morphological details. Second, generative models occasionally produce erroneous predictions with low fidelity to the corresponding true organelle fluorescence image, namely omissions, hallucinations, and structural deformations. These limitations restrict the use of in silico labeling to applications that are not sensitive to fine details or to applications that can cope with a large amount of noise, such as nuclei segmentation (for tracking ^11,12^), and high-content image-based screening ^13^.

Diffusion models (DMs) iteratively refine noise into high-fidelity images ^14,15^. They are trained to reverse the gradual addition of small amounts of noise by estimating the noise that was computationally added (“diffused”) at each step. This training mechanism leads to a model that can generalize well across various noise levels by predicting the noise that needs to be subtracted to make the image slightly less noisy, moving it one step closer to the original image. At inference, DMs generate an image by starting with a pure random noise image and gradually “denoising” it step by step, a process which avoids the instability of single-step predictions. Although originally developed for unconditioned or text-conditioned image generation, DMs have also been adapted for image-to-image translation tasks. This is achieved by conditioning the model on an input image (or its embedding), guiding the denoising process toward a specific output rather than an arbitrary sample ^16^. Thus, DMs learn how to smoothly denoise images, and by incorporating structural priors they can generate new images that align with a given input. Unlike GANs which directly generate images from a noise distribution, the gradual denoising of DMs enables the generation of more realistic images, as demonstrated in many domains, including medical ^17–19^ and biological ^20–22^ imaging.

Fluorescent images are inherently different from natural images. Fluorescent images are characterized by a higher contrast, highlighting specific labeled structures against a dark background. In addition, fluorescent images present a low signal-to-noise ratio (SNR) due to the weak light signal emitted by the fluorophores. Unlike the additive Gaussian noise typical of natural photography, noise in fluorescence microscopy is dominated by Poisson (shot) noise, which is signal-dependent and fundamentally tied to the discrete nature of photon emission ^23,24^. Consequently, the ‘inherent noise’ of the sample is not a stationary background but is intrinsically entangled with the biological structure itself. At inference, during the earlier steps of the DMs denoising process, the inherent noise in the fluorescent images is negligible relative to the high magnitude of synthetic noise present in the image. However, in the later stages, the inherent fluorescence noise becomes dominant. Because DMs are typically trained on a standard Gaussian diffusion manifold, they lack the specific prior to distinguish between the synthetic noise and the signal-dependent biological noise they encounter in the final steps. We hypothesized that this noise entanglement limits the accuracy of synthetic noise estimation as the denoising process approaches the target image. While DMs are designed to refine high-frequency details at late denoising steps, we reasoned that the model begins to mistake inherent fluorescence noise for true biological structure, optimizing for sensor fluctuations rather than genuine organelle morphology. Consequently, the model effectively overfits to local noise patterns rather than to biological structures in the final steps, leading to a loss of structural fidelity. This challenge, a paradox where more denoising steps lead to lower biological accuracy, could hinder the full potential of DMs for fluorescence imaging.

Here, we demonstrate that DMs suffer from diminishing performance at the later steps of the denoising process when applied to fluorescence images. To address this limitation, we present the Diffusion Model for In Silico Labeling (DM4ISL), an in silico labeling model that leverages the iterative denoising process and probabilistic properties of DMs to achieve higher fidelity predictions. Instead of relying on a single, potentially noisy point-estimate at the final denoising step, DM4ISL uses a step-wise self-ensemble averaging approach across intermediate denoising steps where the biological signal remains most distinct from the synthetic noise. This strategy effectively marginalizes the stochastic errors introduced by the model’s attempt to fit sensor noise, resulting in predictions with superior structural stability. DM4ISL systematically outperformed alternative architectures for the in silico labeling of six organelles, as verified with comprehensive benchmarking that included pixel-based, distribution-based, and application-appropriate evaluation metrics ^25^. Beyond prediction, the method uses the standard deviation across the intermediate denoising steps to provide an automatic identification of inconsistent regions, enabling detection of erroneous in silico predictions at inference time.

## Results

### DM4ISL: High-fidelity in silico labeling with quantifiable uncertainty through diffusion averaging

We used the public dataset of spinning-disk microscopy paired brightfield-and-fluorescence volumetric (3D) images of genetically edited fluorescent human induced pluripotent stem cell lines (hiPSC) collected and curated by the Allen Institute for Cell Science (Methods) ^26^. We selected six organelles that have been previously in silico labeled using the same datasets ^6,10,27,28^. These organelles include three structures with clear boundaries in the Nucleoli, Nuclear envelope and Nuclei (DNA); and three filamentous structures: Actin filaments (F-actin), Mitochondria and Microtubules.

DM4ISL extends the implementation ^29^ of the original DM architecture ^15^ to enable 3D image-to-image translation by converting 2D neural network layers into 3D layers (Methods). For each organelle we trained a separate DM4ISL model that was optimized to translate the label-free brightfield image to its corresponding fluorescent image. DM4ISL’s basic training and inference adhere to the standard DM methodology (Fig. 1A-B). During training, a random normally distributed noise image, *ε ∼ N(0,1)*, is drawn, and a step *t ∼ U[0,999]* is sampled from a uniform distribution. The experimentally observed fluorescent image *X*_*0*_ and the noise image *ε* are weighted proportionally to *t* (i.e., larger *t* values induce more noise, see Methods) and added to create a noisy fluorescent image *X*_*t*_ (Methods). An attention-based U-Net ^30^ model (termed “denoiser”) is trained to predict ε_Θ_, an estimation of the noise image ε from the noisy fluorescent image *X*_*t*_ , its corresponding label-free brightfield image (*X*_*BF*_), and _*t*_ the step *t* (Fig. 1A). At inference, DM4ISL applies the trained U-Net to iteratively refine a noisy fluorescent prediction 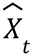 into a slightly less noisy prediction 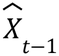 by subtracting a weighted function of the predicted noise image ε_Θ_ (Methods). This iterative denoising process begins with a pure random noise image, 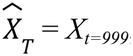, and gradually denoises 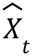, for *t* = *T* − 1,.., 0, until reaching the final fluorescent prediction 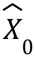 (Fig. 1B).

**Figure 1.**
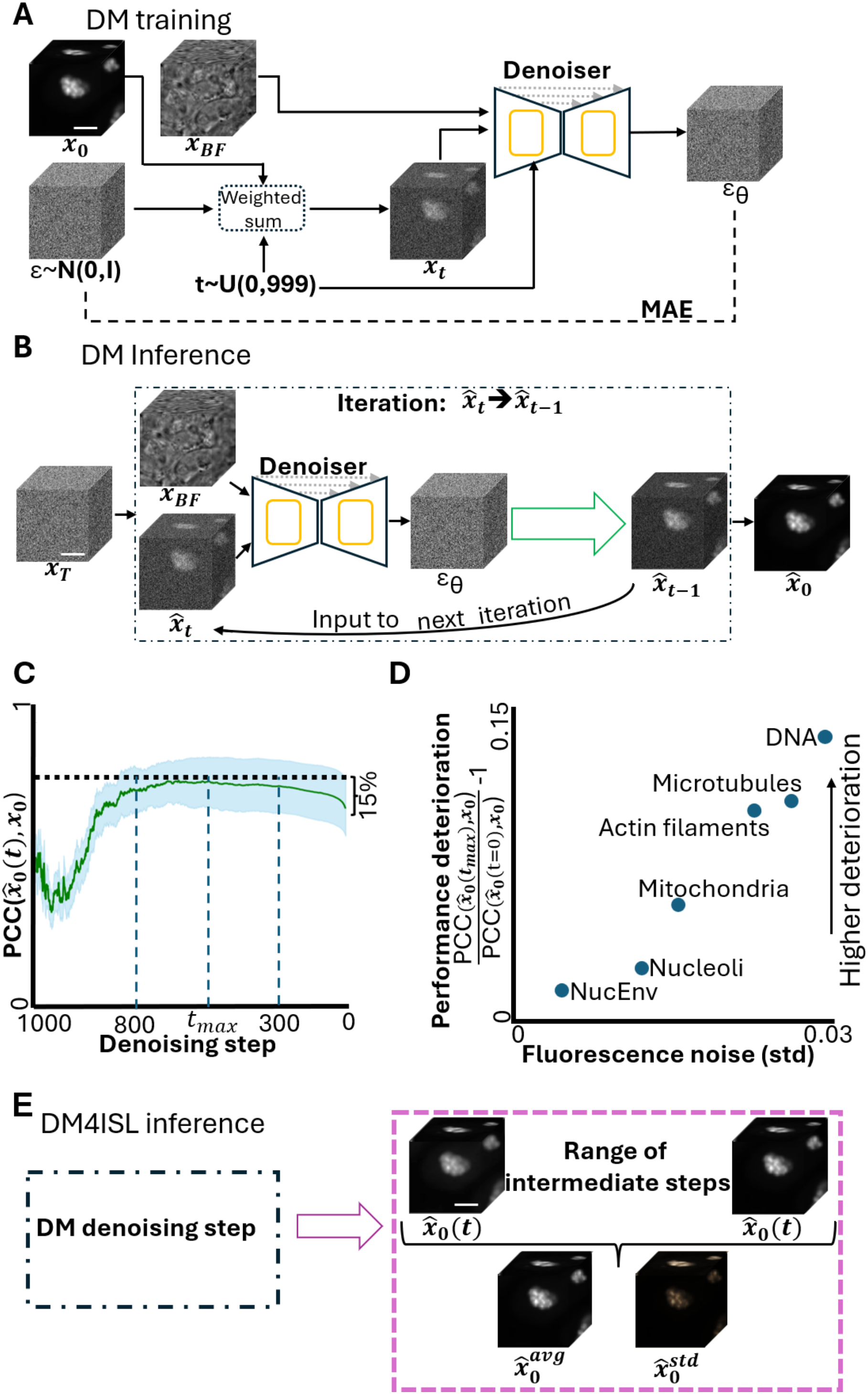
The DM4ISL architecture and its averaging-based inference routine. **(A)** DM training. A noisy fluorescence image, *X*_*t*_ is formed as a weighted sum of the clean image *X*_0_ and Gaussian noise ε∼N(0,I), with weights controlled by a timestep *t* ∼ U(0,999). A U-Net denoiser takes as input the noisy image *X*_*t*_, the corresponding label-free brightfield image *X*_*BF*_, and the timestep t, and predicts ε_Θ_, an estimate of the noise image. This network is optimized using the mean absolute error (MAE) loss between ε and εΘ. Scale bar = 10μm. **(B)** DM inference. A random noise image *X*_*T*_ ∼N(0,I) is denoised by iterative refinement. In each iteration (dashed rectangle), the noisy prediction 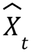 and its corresponding *X*_*BF*_ are provided to the U-Net denoiser, which predicts εΘ. This estimate is then used to produce a less noisy prediction 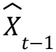 for the next iteration, ultimately yielding the final fluorescence prediction 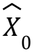. Scale bar = 10μm. **(C)** The evolution of the mean (green) and the standard deviation (cyan) Pearson correlation coefficient (PCC) between the DNA’s fluorescence prediction 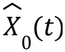 and the corresponding fluorescent images *X*_0_ throughout the iterative denoising process. The horizontal dashed black line depicts the optimal 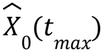, emphasizing a deterioration of 15% in the final fluorescence prediction 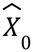. **(D)** In silico labeling performance deterioration (y-axis) as a function of the fluorescence noise (x-axis) for six organelles. For each organelle, the y-axis reports the improvement of the PCC between the optimal 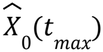 and the ground-truth in respect to the PCC between the final fluorescence prediction 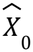 with the ground-truth fluorescence image *X*_0_. A higher ratio indicates greater deterioration in the performance of the final fluorescent prediction 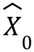 with respect to the optimal prediction 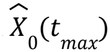. The x-axis shows the standard deviation of the fluorescence noise in void image regions (i.e., without cells) for each organelle. The performance loss was associated with the fluorescence noise across organelles (PCC r = 0.97, p-value < 0.003). **(E)** The DM4ISL component applied at inference. The mean and standard deviation predictions 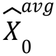 and 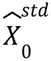 are computed from the fluorescence predictions along the range of intermediate steps, 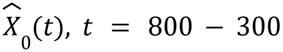 (Methods).

To test our hypothesis that the low SNR of fluorescence images challenges diffusion models at the late stages of denoising, we composed a test set of 128 image patches from the test set’s fields of view that were not used during training (Methods), and evaluated the model’s ability to directly infer the original DNA fluorescent image at each step of the diffusion process.

More specifically, at each denoising step t, we computed 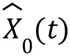, the model’s estimate of the clean image *X*_0_, and evaluated the evolution of these step-wise predictions. At a given step *t*, the noisy fluorescent prediction 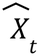 was translated to the final fluorescence prediction 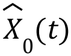 by mapping the current sample directly to the clean manifold using the predicted noise image ε_Θ_ ^31^ (Methods). The evolution of the Pearson correlation coefficient (PCC) between this final fluorescence estimate 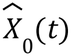 and the ground truth fluorescence image *X*_0_ revealed a pattern (Fig. 1C). At the early steps of the denoising, the DM poorly estimates the noise, leading to large errors in the estimation of *X*_0_. These errors are gradually reduced along the denoising process, as the synthetic noise is progressively removed, until reaching a plateau. In the later denoising steps, the estimation of *X*_0_ gradually declines, revealing an approximate gap of 15% between the maximal PCC observed during denoising and the PCC of the final prediction. A similar trend of deteriorated performance in the late stages of denoising appeared for all five additional organelles, ranging from 2% for the nuclear envelope to 13% for Microtubules (Fig. S1). To verify the correlation between the noise in the fluorescence image and the deterioration of the in silico labeling in the later denoising steps, we estimated the fluorescence noise of each organelle using void (i.e., without cells) regions in the corresponding fluorescence images (Methods). The decline in performance at the late stages of denoising (Fig. 1C and Fig. S1) was associated with the fluorescence noise (Fig. 1D), suggesting that higher fluorescence noise leads to greater degradation in the final in silico labeling.

To further confirm that the diminished in silico labeling performance during the late denoising steps was due to noise in the fluorescence images, we evaluated a diffusion model’s performance on a “super-resolution” reconstruction task, translating low-resolution fluorescence images to high-resolution ones ^32^. Specifically, we downsampled the fluorescence images from the Allen Institutes’ DNA dataset and trained a DM to reconstruct the original high resolution image (Fig. S2A, Methods). This model also showed declined image reconstruction accuracy at the late denoising steps (Fig. S2B). Finally, we evaluated the task of translating Hematoxylin and Eosin (H&E) images to their corresponding DAPI immunofluorescence images, both acquired from the same tissue sections ^33^. In this dataset, the DAPI images were preprocessed to remove noise, thus creating a near noise-free fluorescence target. As expected, the DM was able to virtually stain the H&E to the corresponding DAPI label (Fig. S2C) without exhibiting the late-stage performance decline seen in fluorescence prediction (Fig. S2D). Synthetic addition of noise to the DAPI channel (Methods) led to declined image reconstruction at the late denoising steps (Fig. S2E).

Cumulatively, these results support our hypothesis that the DM struggles to distinguish the computationally synthesized noise from the inherent noise in the fluorescence distribution, leading to deterioration in the final in silico labeling prediction and in other applications of image-to-fluorescence image translation.

In light of this observation, we introduce an inference-time aggregation step, computing the averaged prediction 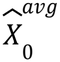 and the corresponding standard deviation 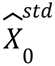, across the fluorescence predictions 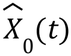 derived from the intermediate denoising steps (Fig. 1E). The range of intermediate steps was empirically set to *t* = 800-300 based on the DNA channel (Fig. 1C; Methods) and was validated for the other organelles and for the application of super-resolution. This range was robust (Table S3) captures the phase where the model resolves the coarse and fine biological structures before diverging due to the high-frequency sensor noise. The averaged prediction 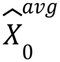 was set as the final in silico labeling prediction, while the standard deviation 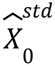 provided a spatial estimate of prediction stability across the denoising process. We call this new model “Diffusion Model for In Silico Labeling” (DM4ISL), providing both a high-fidelity prediction and a corresponding uncertainty map for each inference.

### Qualitative assessment of diffusion-based models for high-fidelity in silico labeling

We compared the in silico predictions of DM4ISL versus two alternative common architectures for cross-modality image translation: A regression network Attention U-Net ^30^, and a generative adversarial network (GAN) Pix2Pix ^34^ (Fig. S3, Methods), as well as the standard DM’s final prediction (Fig. 1B). To allow a fair comparison we used the same Attention U-Net denoiser architecture and applied the same data normalization, data augmentation, batch size and optimization scheme across all models (Methods). Qualitative assessment of the models’ predictions showed that all models were successful in capturing the organelles’ localization, their coarse shape and structure (Fig. 2). However, the diffusion models’ in silico predictions appeared more realistic and crisp than those produced by the Attention U-Net and the Pix2Pix GAN models. Both baseline models tended to produce oversmoothed predictions. For the Pix2Pix GAN, this may reflect the challenge of optimizing adversarial loss across complex 3D biological volumes, thereby losing fine detailed structures. These observations were visualized at the in-focus axial focal plane (Fig. 2) and verified in 3D (Fig. S4). For example, the bright spots in the DNA channel, that are called puncta or foci ^35^, were blurred in the in silico fluorescence images generated by the Attention U-Net and the Pix2Pix GAN. In contrast, the diffusion models generated sharp, clear, and well-defined puncta structures (Fig. 2A). In another example, the Nuclear envelope of a cell undergoing mitosis appeared heavily smudged in the predictions of the alternative approaches, whereas the diffusion models produced sharper contours that more closely resembled the ground-truth fluorescence image (Fig. 2B). Similarly, the internal surface texture of the Nucleoli was smoothed out by the other models, while the DM variants were able to reconstruct some of the fine details (Fig. 2C). The Actin filaments and the Microtubules were extensively smeared by the non-diffusion models, whereas the diffusion models generated images that better resembled the true structure (Fig. 2D-E). Our visual assessment of the Mitochondria did not identify clear benefits of one model over the others (Fig. 2F). While both diffusion-based approaches provided superior visual clarity over

**Figure 2.**
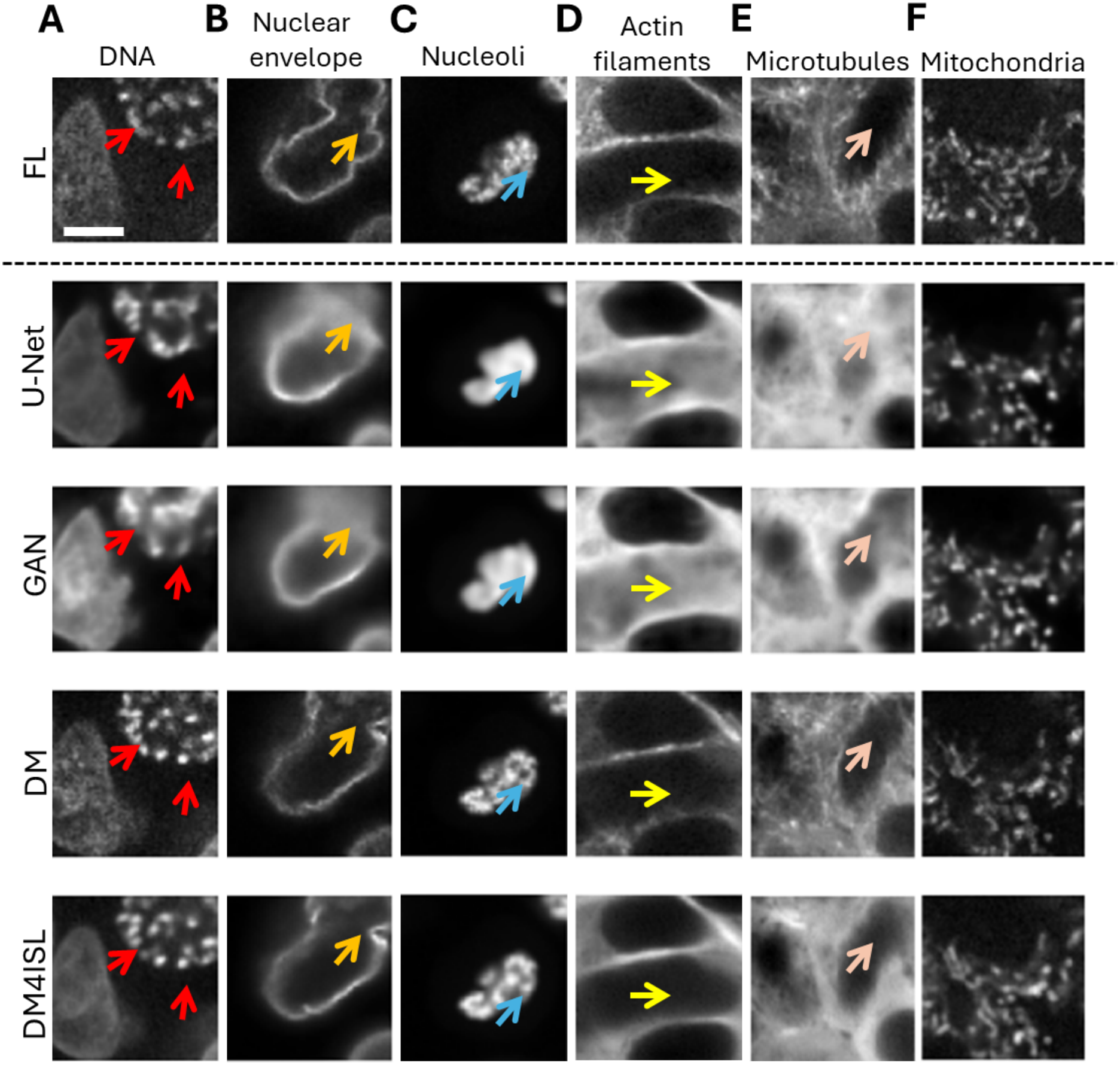
Visual qualitative assessment of in silico labeling models. The first row shows one representative fluorescence image for each of the six organelles. The other rows display the corresponding in silico labeling of (top-to-bottom): U-Net, GAN, DM and DM4ISL models. Colored arrows guide to regions where the Attention U-Net and Pix2Pix GAN models deviate from the fluorescence image. **(A)** Blurred puncta in the DNA channel (red arrows). **(B)** Smudged nuclear envelope for a cell undergoing mitosis (orange arrow). **(C)** Smoothed nucleoli surface (cyan arrow). **(D)** Smeared Actin filaments (yellow arrow). **(E)** Smeared Microtubules (pink arrow). **(F)** No major deviations were observed for the Mitochondria. Scale bar = 10μm.

U-Nets and GANs, direct qualitative comparison between the standard DM and DM4ISL did not reveal a clear visual advantage for either method. This underscores the necessity for the rigorous quantitative benchmarking presented in the following section to resolve the subtle but critical differences in structural fidelity. Altogether, these qualitative visual assessments indicate that diffusion models generate predictions that more faithfully represent fine subcellular structures compared to common non-diffusion architectures.

### Quantitative assessment of DM4ISL for in silico labeling

We composed a test set for each organelle consisting of 128 image patches randomly selected from 8-19 fields of view not seen during training. We devised three sets of image quality metrics to evaluate the similarity between the in silico predictions and their corresponding ground-truth fluorescence images. Specifically, for each organelle and for each metric, we calculated a similarity score for every image patch and reported the mean matched performance across the entire test set. The first set of metrics calculate similarities at the pixel and localized patch-scales: (1) pixelwise mean-square error (MSE); (2) multiscale structural similarity index measure (MS-SSIM), a weighted average of the structural similarity index (SSIM) that considers changes in structural information, brightness, and contrast, across multiple downsampled scales ^36^ and (3) image patch correlation using Pearson’s correlation coefficient (PCC). The second set of metrics calculate similarities between image-based intensity distributions: (4) mutual information (MI) that quantifies the statistical dependency between the joint intensity distribution of two images and their individual intensity distributions ^37^. Ultimately, in silico labeling predictions should be validated by assessing their practical value in enabling specific downstream biological analyses, such as organelle localization, counting, and shape analysis ^5,25,38^. This approach helps determine the magnitude of the biological signal that is required to make a concrete biologically interpretable conclusion. Moreover, improvements in image quality metrics do not always translate to improved performance in downstream analysis tasks ^39^. Thus, the third set of metrics includes six “application-appropriate” measurements ^25^ according to downstream analysis tasks that require organelle segmentation. One application-appropriate measurement is (5) Intersection over Union (IoU) of the semantic organelle segmentation masks. This metric requires a classical semantic segmentation pipeline that in our case included filtering, thresholding and removal of small objects (Methods). The remaining five metrics rely on 2D watershed-based instance segmentation ^8,13,40^ and were applied to structures with distinct physical boundaries: DNA, Nuclear envelope and Nucleoli (Methods): (6) Instance F1 score (F1) evaluates detection accuracy to pair predicted and ground-truth instances; (7) Centroid distance (CD) measures spatial displacement (in *µ*m) between paired centroids; (8) Nearest-neighbor distance error (ND) assesses the spatial relations between adjacent instances by calculating the distance (in *µ*m) between their contours; (9) Area discrepancy (Area) measures the size error between paired instances (in *µm*^2^); (10) Hausdorff distance (HD) measures the maximal boundary alignment error between paired instances (in *µ*m) ^41^.

We benchmarked the four in silico models over these metrics. DM4ISL outperformed the U-Net and the GAN models in 41 of the 45 (∼91%) organelle-metric pairs, with an average improvement of 15.8% and 20.1% respectively across all metrics and organelles (Fig. 3A-F). While DM4ISL did not show a qualitative advantage over DM, our quantitative benchmarking showed a consistent superiority of DM4ISL in 37 of the 45 organelle-metric pairs, statistically significantly worse only for the Actin filament MSE. Overall, DM4ISL achieved an overall average improvement of 8.5% across all metrics and organelles compared to DM. This advantage was most pronounced for the Mitochondria and the DNA, with an average improvement of 23.3% and 15.4%, over all metrics, correspondingly. Nuclear envelope and Nucleoli were the organelles that were least improved by DM4ISL with an average gain of 4% and 5% respectively over all metrics. Interestingly, actin filaments showed a 37.8% deterioration in MSE for DM4ISL relative to standard DM, thus suggesting that while DM4ISL better reconstructs the statistical information of actin filaments, it struggles with the absolute pixel-wise error. Focusing on application-appropriate measurements, the most relevant metrics in terms of their biological interpretation, DM4ISL showed a consistent average gain of 3-8.6% for each of the six metrics and across all organelles, compared to DM. The full results are available in Tables S1-S2. These results confirmed that by avoiding the noise-entangled final steps, DM4ISL captures a more stable biological signal than the standard DM final-step inference.

**Figure 3.**
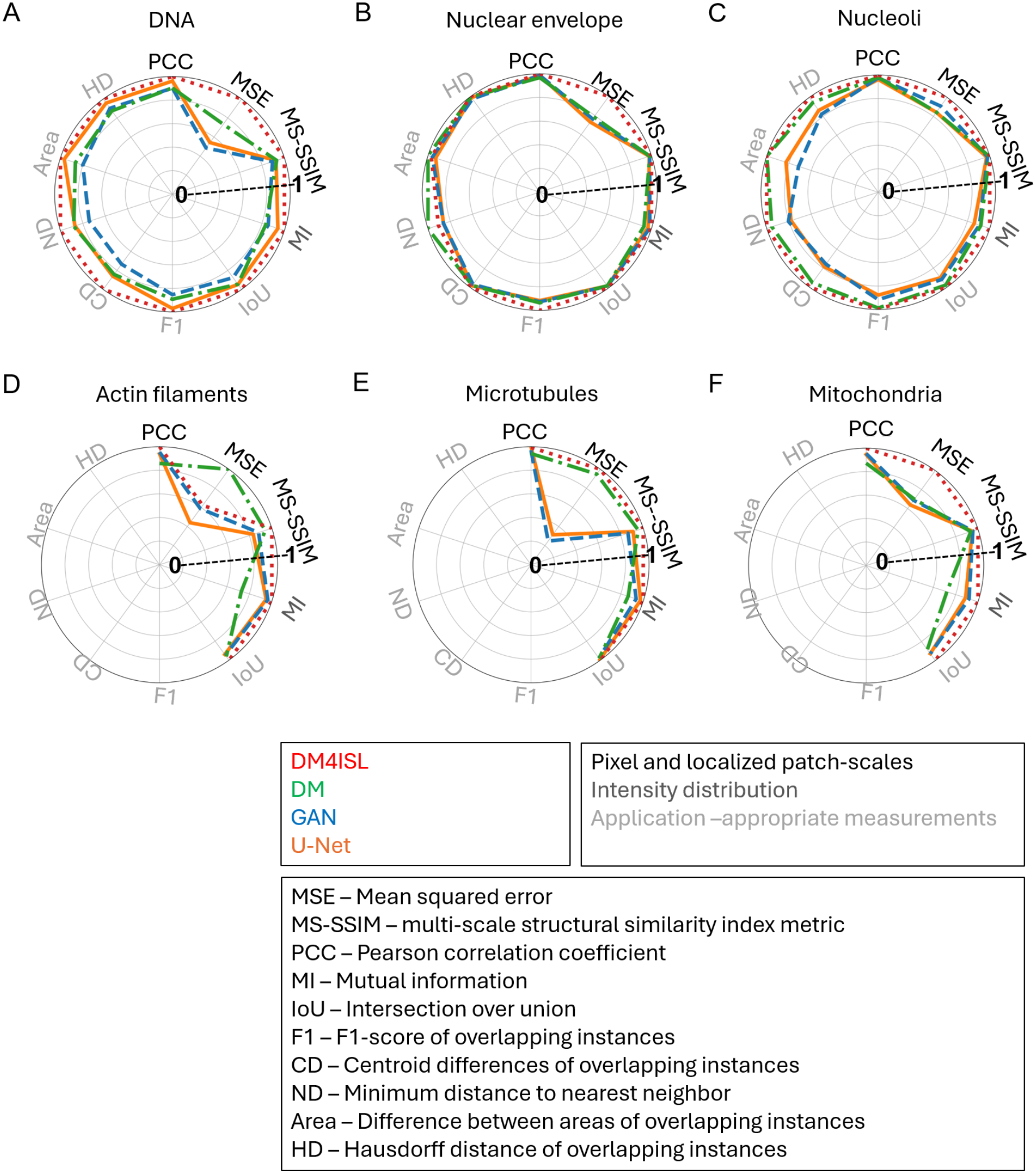
Benchmarking in silico labeling across different models and metrics. Benchmarking the Attention U-Net (orange), Pix2Pix GAN(blue), DM (green), DM4ISL (red), using the per-organelle test set of N = 8-19 fields of view with n = 128 randomly selected image patches. Ten metrics are displayed in six polar plots, one per organelle. Each metric was scaled by setting the best model to a value of 1 (Methods). Three types of image quality evaluation metrics: pixel and localized patch-scales (black), intensity distribution (dark gray), and application-appropriate measurements (light gray). Full details on the evaluation metrics are available in the Methods. Full results and statistical analyses are available in Table S1 and Table S2. **(A)** DNA. **(B)** Nuclear envelope. **(C)** Nucleoli. **(D)** Actin filaments. **(E)** Microtubules. **(F)** Mitochondria.

### Uncertainty estimation with DM4ISL detects erroneous predictions at inference

The utility of in silico labeling is constrained by the propensity of deep learning models to generate structural errors that propagate through downstream analysis, leading to biased biological quantifications ^42^. These errors typically manifest in three forms: deformations, where the predicted morphology is distorted, omissions, where existing biological structures are omitted, and hallucinations, where the model generates plausible-looking structures absent from the ground truth. Without a mechanism to identify these failures at inference time, such inconsistencies remain indistinguishable from high-fidelity predictions.

To assess the susceptibility of DM4ISL to generate structural errors, we visually inspected all 128 DNA image patches from the test set and manually annotated those that exhibited unambiguous substantial errors in their in silico labeling predictions for DM4ISL and for the Attention U-Net as a baseline, where an “unambiguous substantial error” was defined as a visually apparent structural deformation, omission, or hallucination. According to this analysis, many test image patches contained substantial errors in their U-Net in silico labeling predictions (Fig. 4A). Systematic manual quantification revealed that 44 out of the 128 (∼34%) image patches contained unambiguous substantial in silico labeling errors, which were dominated by deformations (Fig. 4B). In contrast, DM4ISL exhibited unambiguous substantial in silico labeling errors in 25 of the 128 (∼20%) DNA image patches (Fig. 4C).

**Figure 4.**
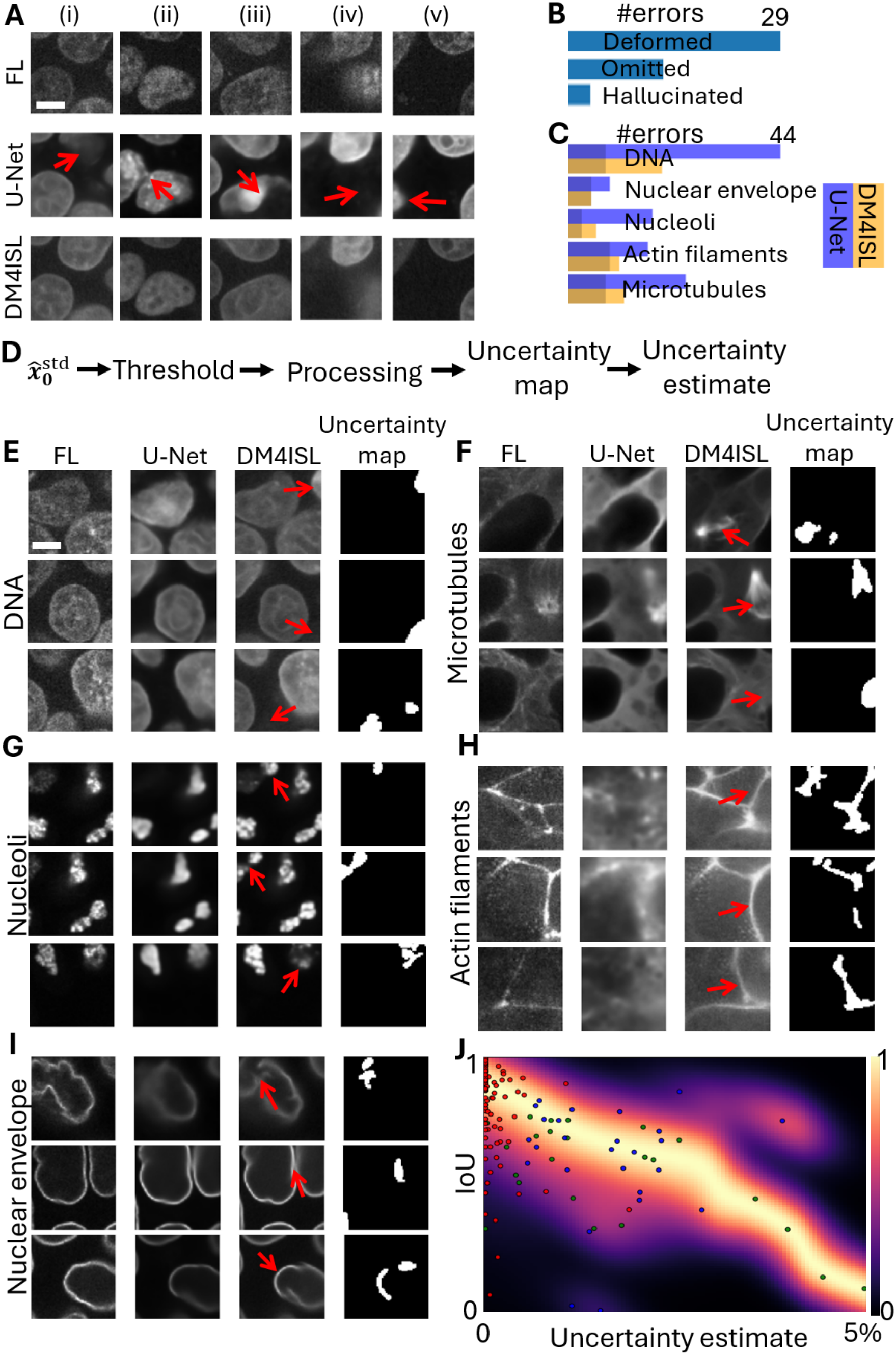
DM4ISL can detect erroneous predictions at inference. **(A)** Representative erroneous predictions of the DNA (top-to-bottom): fluorescence image patches (‘FL’); the corresponding Attention U-Net’s (‘U-Net’) in silico labeling; and the corresponding DM4ISL’s in silico labeling. Red arrows guide to regions where the U-Net substantially deviates from the fluorescence image (columns): (i-iii) Deformed nuclei; (iv) Omitted nucleus; (v) Hallucinated nucleus. Scale bar = 10μm. **(B)** Quantification of manually annotated unambiguous substantial errors for in silico labeling of the DNA by the U-Net model. Overall, 44 out of the 128 DNA image patches contained errors in their U-Net predictions. These were partitioned into 29 deformations, 12 omissions and 3 hallucinations. **(C)** Manually annotated unambiguous substantial errors for in silico labeling by the Attention U-Net (purple) and DM4ISL (orange) for five organelles: DNA (44 versus 25 errors), Nuclear envelope (9 versus 5 errors), Nucleoli (18 versus 6 errors), Actin filaments (18 versus 11 errors) and Microtubules (25 versus 12 errors). Opaque regions represent erroneous predictions of instances shared by both models. Representative erroneous U-Net predictions of the Nuclear envelope, Nucleoli, Actin filaments and Microtubules are displayed in Fig. S5. Erroneous predictions in Mitochondria were too difficult to assess by eye and thus were not included in this analysis. **(D)** Image analysis pipeline for generating the binary ‘uncertainty map’ from DM4ISL’s 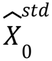 (**Methods**). **(E-I)** Representative DM4ISL-derived uncertainty maps flagging manually annotated substantial erroneous DM4ISL predictions. Each panel represents an organelle, displaying three instances (rows), and including (left-to-right): the fluorescence image patch (FL), the corresponding U-Net fluorescence prediction (U-Net), the corresponding DM4ISL fluorescence prediction (DM4ISL), and the corresponding uncertainty map. Red arrows guide to regions of the substantial erroneous DM4ISL predictions. **(E)** DNA. **(F)** Microtubules. **(G)** Nucleoli. **(H)** Actin filaments. **(I)** Nuclear envelope. Scale bar = 10μm. **(J)** Density Resampled estimate of Mutual Information ^43^ between the DM4ISL-derived uncertainty estimate and the intersection-over-union (IoU) measurement for the DM4ISL’s DNA predictions. DREMI estimates the conditional density P(Y|X), resamples evenly across X to remove sampling bias. The N = 128 image patches’ values are overlaid on top of the DREMI plot, with Pearson’s correlation coefficient of -0.5 (p-value < 0.03) between the uncertainty map and the IoU of these patches. Green indicates manually annotated substantial errors correctly flagged by the uncertainty estimate; Blue indicates prediction errors that affected downstream analysis tasks, but were too subtle for the human eye; Red are the remaining image patches.

Expanding this manual quantification to additional organelles confirmed that DM4ISL was less prone to unambiguous substantial errors, with 59 error-containing patches (∼9%) compared to 114 for the U-Net predictions (∼18%) across the 640 image patches of the five assessable organelles (Fig. 4C). Mitochondrial predictions were excluded from this manual analysis as their high structural complexity and density made erroneous predictions too difficult to reliably assess by eye. Interestingly, manual comparison of DM versus DM4ISL showed that both models exhibited unambiguous substantial in silico labeling errors in the same image patches, indicating that the benefit of DM4ISL stems from higher-fidelity reconstruction of fine structural textures and boundaries rather than the elimination of major structural artifacts, which are shared across both diffusion-based approaches. Altogether, these results establish that diffusion models are less susceptible to substantial in silico labeling errors, as defined by a human expert, compared to the U-Net architecture. By design, DM4ISL provides a dual output at inference: a high-fidelity averaged prediction 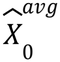 and a corresponding spatial standard deviation map 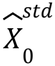 that reflects the local prediction stability.

Having established that diffusion-based models reduce, but do not entirely eliminate, substantial in silico labeling errors, we sought to determine if this internal standard deviation could serve as a reliable diagnostic tool. We hypothesized that regions of high 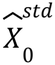 would highlight areas where the model’s denoising trajectory is unstable, thereby flagging localized errors at inference. To test our hypothesis, we generated an ‘uncertainty map’ by applying an organelle-specific threshold to 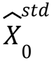, followed by applying morphological operations to discard isolated, small-scale noise (Fig. 4D, Methods). The fraction of positive voxels served as an “uncertainty estimate” to flag regions of low prediction stability (Methods). This uncertainty estimate reached high sensitivity for the manually annotated substantial errors across five organelles (excluding Mitochondria). All but four of the 59 erroneous manual predictions were successfully flagged by the uncertainty map (Fig. 4E-I, Fig. S6).

Furthermore, the uncertainty estimate flagged 16 errors across organelles that were overlooked during the manual inspection (Fig. S7). Examining false alarms (false-positives) flagged by the uncertainty estimate revealed 26 DNA image patches that were not identified during manual inspection because they lacked obvious visual artifacts. Closer examination revealed that these false alarms were mostly attributed to abnormal cell shapes (Fig. S8).

Systematic analysis of the DNA patches confirmed that the uncertainty estimates were associated with their corresponding “application-appropriate” IoU measurements (Methods, Fig. 4J). Overall analysis demonstrated high error detection performance, correctly identifying 55 true positives (93.2% sensitivity) and 537 true negatives (95.0% specificity), against 4 false negatives (6.8% false-negative rate) and 28 false positives (5.0% false-positive rate, of which 26 corresponded to dysmorphic DNA cell shapes). These results indicate that DM4ISL’s uncertainty estimation provides a built-in mechanism enabling automated identification of erroneous predictions at inference.

## Discussion

The field of bioimage informatics is founded on the fundamental premise that biological images exhibit unique physical and statistical properties that distinguish them from natural images ^25,44,45^. Fluorescence microscopy, in particular, is governed by specific constraints, such as Poisson-limited photon noise, fluorescence decay, and the diffraction-limited nature of subcellular structures. Despite this theoretical distinction, the practical evolution of the field has largely been driven by training or fine-tuning “off-the-shelf” architectures designated for natural images under the assumption that they are robust enough to generalize across domains ^46^. Explicit examples where biological signal properties actively hamper the performance of standard architecture remain surprisingly rare. While the community has documented general pitfalls such as the pixel-wise blurring inherent to MSE-optimized U-Nets or the stochastic hallucinations produced by standard GANs ^47^, these are often viewed as unavoidable trade-offs rather than specific failures of the underlying generative logic ^48^.

Our study reveals that the standard late-stage denoising trajectory of diffusion models, while optimized for high-frequency detail in natural scenes, becomes counterproductive for fluorescence in silico labeling. In this specific context, the final steps of the diffusion process do not “refine” the true biological signal but rather entangle the inherent noise in the fluorescent images with the synthetic noise because, at later stages of the diffusion process, these two factors become challenging to distinguish for a standard denoiser. By introducing DM4ISL and its intermediate prediction averaging, we provide a domain-specific refinement that mitigates this conflict, distilling the biological signal over the generative noise.

Beyond theoretical advantages, our qualitative and quantitative assessments demonstrate that diffusion-based architectures, and particularly DM4ISL, consistently outperform alternative models in structural fidelity and predictive reliability across organelles. By employing a suite of metrics and especially application-appropriate metrics ^25^, we reported DM4ISL’s enhanced capacity for the reconstruction of fine structural textures and boundaries that were frequently lost by the alternative models. These results align with previous studies reporting enhanced performance of diffusion models in medical ^17^ and biological ^20–22,49–51^ imaging. Crucially, our manual quantification revealed that diffusion models reduced the incidence of unambiguous substantial errors by almost 2-fold compared to the U-Net baseline. Notably, both the standard DM and DM4ISL exhibited a similar reduction in these major artifacts, suggesting that the iterative denoising process itself is more robust against the shape deformations and structural mergers seen in traditional image-to-image translation models. However, while the diffusion framework reduces major errors, DM4ISL’s intermediate prediction averaging provides the additional accuracy required to marginalize subtle high-frequency fluctuations that translate to better performance in downstream applications. Consequently, DM4ISL yields in silico labels that are not only visually superior but demonstrate the structural stability necessary for rigorous biological quantification.

The ability to identify where a model fails is critical toward using in silico labeling for biological discovery ^52–54^. DM4ISL addresses this challenge by leveraging the statistical standard deviation across intermediate denoising steps to generate a localized uncertainty map. This map serves as a built-in diagnostic tool, quantifying the stability of the generative process under the assumption that non-stable regions are associated with prediction errors ^55^. Importantly, our approach provides this uncertainty in a single inference trajectory, without requiring an ensemble of models ^56–58^, multiple inference passes ^32,59,60^, or alteration of the training process ^52–55^. The highly accurate empirical identification of manually annotated localized substantial errors, along with the capacity to flag structural discrepancies that initially escaped expert human inspection, provides a robust method for trustworthy in silico labeling. While DM4ISL is currently limited by the computational complexity inherent to iterative denoising ^15^ and the requirement for organelle-specific parameter tuning, future iterations using latent diffusion ^61^ or flow-matching ^58,62^ architectures may improve inference speed. Ultimately, this automated, data-driven error detection acts as a rigorous quality control layer, ensuring that in silico predictions can move beyond qualitative visualization toward reliable, large-scale biological quantification.

### Declaration of generative AI and AI-assisted technologies in the writing process

During the preparation of this work the authors used Gemini and ChatGPT in order to edit and refine the text. After using this service, the authors reviewed and edited the content as needed and take full responsibility for the content of the manuscript.

## Methods

### Dataset and pre-processing

We utilized the public dataset of 3D spinning-disk confocal microscopy images of genetically edited fluorescent human induced pluripotent stem cell (hiPSC) lines from the Allen Institute for Cell Science ^26^. We focused on six organelles, three of which with clear boundaries (Nucleoli, Nuclear envelope, DNA), and three with filamentous structures (Actin filaments, Mitochondria, and Microtubules). Following ^6^, the 3D field of view images were resized to 366×244 pixels in the x-y plane to achieve a cubic voxel (0.29 *µ*m). Each field of view was cropped into 64×64×16 pixels image patches. Each image patch was normalized to the range [0,1] by first clipping intensities to a fixed organelle-specific lower and upper bounds, followed by min-max scaling using these bounds, for both brightfield and fluorescence data. To train and evaluate the in silico labeling models on image patches containing cells, rather than patches dominated by void or out-of-focus, we excluded patches according to their fluorescence pixel-intensity standard deviation (std) at the slice-level. Specifically, after intensity clipping and prior to min-max scaling, using ∼5000 randomly selected training patches, we manually determined organelle-specific std thresholds separating between void or out-of-focus and fine organelle structure: Nuclear envelope (70), Nucleoli (53), Mitochondria (18), Actin filament (25), Microtubules (250), and DNA (20). A patch was retained only if its fluorescence intensity std exceeded the corresponding threshold in every XY slice. The lower threshold for DNA reflected its abnormally low signal-to-noise ratio, having higher background noise relative to all other organelles (Fig. 1D), while also exhibiting relatively low standard deviation in regions containing foreground and background. The threshold was therefore set to ensure that dim or structurally variable cellular regions were retained. Consequently, this inclusive threshold retained a broader diversity of patches, including those containing dysmorphic cells or intense local foreground clusters (Fig. S8).

### Diffusion model architecture

To perform in silico labeling of volumetric fluorescence images from label-free brightfield (BF) data, we implemented a three-dimensional (3D) Denoising Diffusion Probabilistic Model ^15^ (DM). The model was designed to learn the target distribution of organelle-specific fluorescence by reversing a forward diffusion process. In the forward process, a ground truth fluorescence image *X*_0_ was corrupted by additive Gaussian noise *ε*∼N(0,1) according to a variance scheduler α_*t*_ over T=1000 discrete timesteps. The resulting noisy image *X*_*t*_ at any step [0,T-1] was defined as:

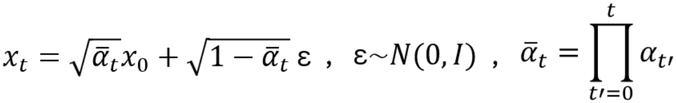

We utilized a noise scheduler α which evolved from 0.9995 to 0.9805:

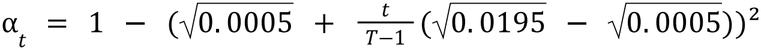

The backbone of our model was an Attention U-Net architecture implemented via the MONAI framework ^29^. The architecture comprises a contracting encoder and an expanding decoder with four spatial scales (input size 64×64×16 voxels). The architecture was adapted from 2D to 3D operators to handle the volumetric input by modifying the convolutional layers to 3D. The network consists of four downsampling and upsampling levels with channel depths of [128, 256, 256, 512] corresponding to three downsampling and upsampling operations. To capture long-range spatial dependencies while managing computational overhead, self-attention modules were integrated at the bottleneck layer, the deepest layer of the U-Net, where the spatial resolution is lowest (8×8×2 voxels). Training was conducted using the Adam optimizer with a learning rate of 2.5e-5 and a batch size of 16 patches.

Models were trained for 100K iterations.

The Attention U-Net denoiser network learns the Gaussian noise used to corrupt the fluorescence image *X*_0_, while conditioning on the label-free brightfield image patches, *X*_*BF*_, by concatenating *X*_*BF*_ and *X*_*t*_ along the channel dimension. The model was optimized by minimizing the mean absolute error (MAE) between the noise *ε* and the predicted noise *ε*_θ_:

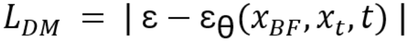

During inference, a random noise volume *X*_*T*_ ∼*N*(0, 1) is iteratively denoised for t = T-1, …,0. At each step, the model predicts the noise component *ε*_θ_, allowing for the calculation of the refined volume 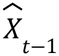 using a stochastic transition:

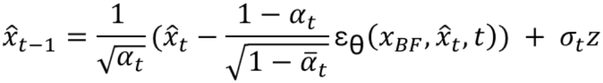

where *z*∼*N*(0, 1) and *σ*_t_ is defined by the noise schedule ^15^:

### DM4ISL

DM4ISL was defined as the ensemble average 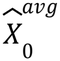 of the estimated clean image, 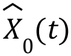, across a stable intermediate steps interval, where

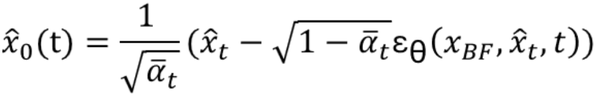

and

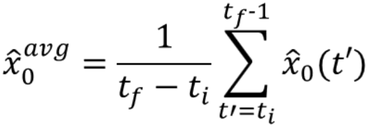

The averaged predictions were subsequently min-max normalized to [0,1] across all patches processed in the same inference call. The interval of the range of intermediate denoising steps was empirically set to *t*_*i*_ = 300 *to t*_*f*_ = 800 for all organelles. We tested the sensitivity of the range by calculating the Pearson correlation coefficient for several other endpoints around the selected range for a subset of samples from the test data (Table S3).

The standard deviation 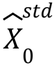 across the interval of intermediate denoising steps was used to provide a spatial estimate:

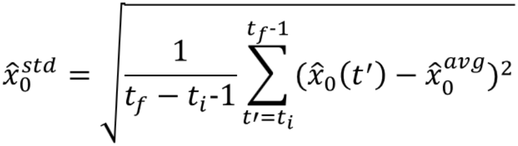

### Estimating the diffusion model’s deterioration along the diffusion process

We estimated the performance deterioration along the denoising process of the DM by comparing the maximum PCC value at intermediate steps and the PCC value at the final step (Fig. 1C-D):

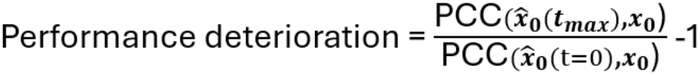

Thus, a deterioration value of 15% indicates that the maximum intermediate PCC was 15% higher than the final PCC.

### Estimating the fluorescence noise

We estimated the fluorescence noise (Fig. 1D) of each organelle by calculating the standard deviation (std) across ∼20 void fluorescence patches (i.e, without cells) selected manually out of randomly drawn patches from the fields of view.

### Train-test data partitioning and data augmentation

To avoid data leakage we partitioned train-and-test sets at the FOV level. For each organelle, 90% of the FOVs were assigned to the training set, while the remaining 10% (8–19 FOVs per organelle) were reserved as a held-out test set. Performance was evaluated on 128 patches randomly sampled from the test FOVs. Patches in the training set underwent stochastic data augmentation, including horizontal and vertical flips and discrete rotations of 90°, 180°, and 270°.

### Fluorescence super-resolution reconstruction

Ground-truth DNA fluorescence volumes from the Allen Institute for Cell Science ^26^ were subjected to a 512-fold volumetric downsampling (a factor of 8 reduction in each spatial dimension). These volumes were then linearly upsampled to their original dimensions to generate a degraded input representation. This degraded signal served as the conditional input for the DM, testing its ability to recover high-frequency spatial information from low-resolution fluorescence data. Train-test data partitioning and data augmentation were performed similarly to the application of in silico labeling.

### Virtual staining of histological sections to DAPI channel

Virtual staining of Hematoxylin and Eosin (H&E) to their corresponding DAPI immunofluorescence images was demonstrated using the HEMIT dataset ^33^. These data consisted of 1,239 registered image pairs, of size 1024×1024 pixels each, from the same tissue sections. The dataset was split into training and test sets in a 9:1 ratio. To maintain architectural consistency with the volumetric DM, images were processed as 64×64×1 image patches. To investigate the impact of sensor noise on the diffusion trajectory, we generated synthetically degraded DAPI images by adding a combination of Gaussian noise (variance σ^2^ = 5 intensity units^2^) and Poisson noise to the fluorescence images prior to patch extraction and training.

### Comparative baseline architectures

We benchmarked DM4ISL against in-house implementations of two common architectures for cross-modality image translation:

#### Attention U-Net ^30^

A standard deterministic model utilizing the same encoder-decoder configuration and hyperparameters as the DM backbone. This model was trained to optimize an MAE loss function between the fluorescence image patches *x* and the in silico labeling *x*:

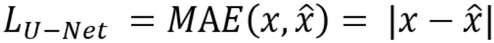

#### Pix2Pix GAN ^34^

A conditional generative adversarial network (cGAN). The generator architecture matched our Attention U-Net. A PatchGAN discriminator was employed to evaluate 4×4×4 local patches, enforcing high-frequency structural realism. The generator was optimized using a composite loss function:

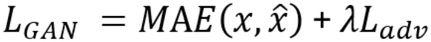

where *L*_*adv*_ is the binary cross-entropy adversarial loss and λ was empirically set to 0.02 to ensure training convergence.

### Quantitative assessment of in silico labeling performance

To evaluate prediction fidelity and biological relevance, we benchmarked all models across ten image quality metrics partitioned into three distinct sets. For each metric, scores were calculated across matched pairs of test-set ground-truth fluorescence patches (*x*) and in silico predictions 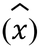, with the mean performance reported across the entire test set on patches with valid scores for all four models.

The first set of image quality metrics consisted of pixel-level and local patch-scale metrics:

(1) Mean Squared Error (“MSE”) is a common metric used to measure the average pixel-wise squared difference between the fluorescence image patch *x* and its corresponding in silico labeling 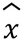:

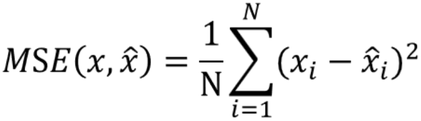

Where N is the number of voxels in the image patch

(2) Multi-scale Structural Similarity Index (“MS-SSIM”). SSIM evaluates perceptual similarity between the fluorescence image patch *x* and its corresponding in silico labeling 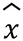 across luminance (l), contrast (c), and structural (s) terms ^36^. MS-SSIM was computed over the XY dimensions of the 3D image patches and extends SSIM by combining the similarity components across multiple downsampled scales, M = 5, using scale specific weights.

Where α, β, γ are the weights assigned for every term across the scales ^36^:

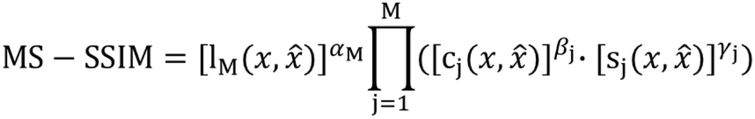

(3) Pearson Correlation Coefficient (“PCC”) measures the linear correlation between the fluorescence image patch *x* and its corresponding in silico labeling 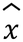:

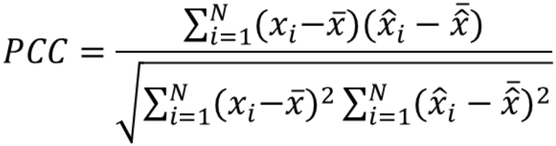

The second set of image quality metrics consists of the information theory metric that characterizes the intensity distributions per patch across all the test set:

(4) Mutual information (“MI”) quantifies the statistical dependency between the joint intensity distribution of the ground-truth fluorescence and corresponding in silico labeling predictions, and their marginal intensity distributions ^37^:

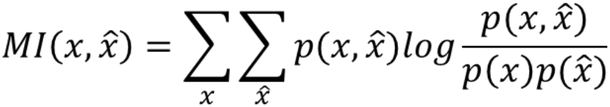

MI was estimated using a joint intensity histogram with 20 bins per image.

The third set of image quality metrics consists of application-appropriate measurements ^25^. These metrics were calculated at the 2D slice level, to refrain from errors which may occur from attempting to connect an instance across 3D slices. In order to evaluate the biological utility of the predictions for downstream analysis, we started by implementing a semantic segmentation pipeline: (i) Gaussian smoothing (sigma=1) to suppress pixel-level noise; (ii) Otsu’s automated thresholding ^64^; (iii) holes filling; (iv) morphological erosion and dilation, using a disk-shaped structuring element with a radius of 1 pixel, to remove background noise segmentation residuals; and (v) removal of foreground objects smaller than 1% of the slice area (41 pixels). This semantic segmentation was used to define the next metric:

(5) Intersection over Union (“IoU”) quantifies the overlap between the semantic segmentation masks obtained from the fluorescence image patch *x* and its corresponding in silico labeling *x*. IoU takes into consideration the foreground true positive (TP) pixels that overlap correctly, false positive (FP) pixels that are predicted as foreground but belong to the background in the fluorescence mask, and false negative (FN) pixels that are foreground in the fluorescence mask but missed in the prediction:

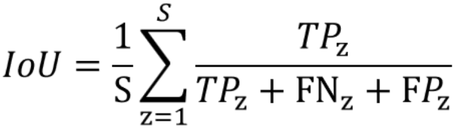

IoU is calculated per slice z and averaged over all S slices of the 3D image patch. Slices in which neither mask contains the foreground are excluded.

The remaining five metrics rely on organelle instance segmentation, and were therefore evaluated on structures with distinct physical boundaries: DNA, Nuclear envelope, and Nucleoli. Filamentous structures (Actin filaments, Mitochondria and Microtubules) were excluded from instance-level evaluation because they cannot be reliably partitioned into discrete instances using standard segmentation, requiring alternative, application-specific metrics.

To generate instance masks, a final watershed transformation ^40^ was applied directly to the binary foreground masks produced by the semantic segmentation stage. Watershed markers were placed at local maxima of the distance transform at least 10 pixels apart, and each connected foreground component was assigned at least one marker. Instances smaller than 1% of the slice size (41 pixels) were then merged into their neighbouring instance by repeating the watershed without their markers. This removes over-segmentation fragments. Using these instance segmentation masks, we calculated five structural metrics ^5,25,38^ comparing ground-truth fluorescent instances with their corresponding in silico labeled predictions. Instance pairs were established per image patch slice via one-to-one Hungarian matching ^65^ based on an Intersection over Union (IoU) cost matrix, retaining only valid matches with an IoU ≥ 0.6 ^66^. Instance-level metrics requiring matched instances were averaged over the matched instances of each slice, then over the slices of each image patch, and finally across all image patches in the test set to yield a single representative score per metric. Slices or image patches with no matched instances do not define these metrics and were excluded from the averages.

(6) Instances F1 score (‘F1’) , evaluating detection and instance segmentation accuracy. Instances from the fluorescent and the in silico labeled predictions with IoU at or above the defined threshold are considered true positives (TP), unmatched fluorescent instances are considered false negatives (FN), and unmatched in silico labeled predictions are considered false positives (FP).

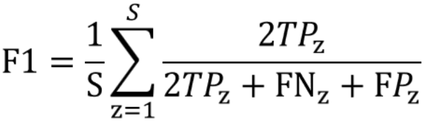

### F1 is calculated over all S slices of the 3D image patch. Slices containing no instances in either image are excluded

(7) Centroid distance (CD) measures the spatial colocalization fidelity.

For matching instances between the fluorescent and in silico labeled images, we calculate the slice-level Euclidean distance between center coordinates:

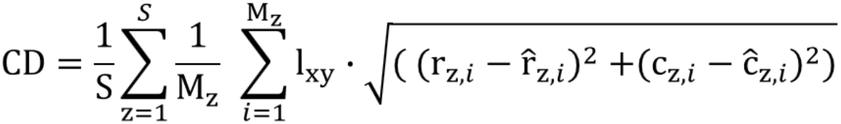

Where (*r*_*z,i*_, *c*_*z,i*_) and 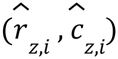 represent the center row and column pixel coordinates for matched instance i in slice z for the fluorescent image patch x and the in silico labeled prediction 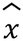. *l* = 0. 29(*µm*/*pixel*). CD (*µm*) is averaged over all matched instances *M*_*z*_ in slice z and then over all valid slices S of the 3D image patch.

(8) Neighbor distance (“*ND*”) evaluates spatial organization and organelle proximity. For each matched instance between the fluorescent and in silico labeled images, when more than one instance is present in both images, we calculate within each image the minimal distance *d*_*min*_ between the instance border and the border of its nearest neighboring instance in the same slice. Touching instances have a distance of 0. ND is calculated as the absolute error between the distances:

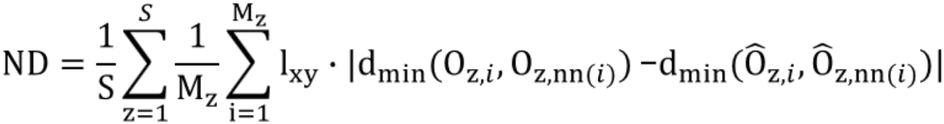

Where *O*_*z,i*_ and *Ô*_*z,i*_ represent matched instance i in slice z for the fluorescent image patch x and the in silico labeled image patch 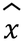, and *O*_*z,nn*(*i*)_ and *Ô*_*z,nn*(*i*)_ represent the nearest neighbors, respectively. *l*_*xy*_ = 0. 29(*µm*/*pixel*). ND (*µm*) is averaged over all matched instances *M*_*z*_ in slice z and then over all valid slices S of the 3D image patch.

(9) Area discrepancy (“Area”) measures the morphological size fidelity.

For matched instances between the fluorescent and in silico labeled images, we calculate the slice-level area absolute error:

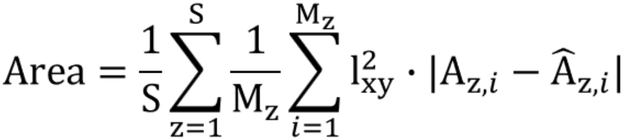

Where *A*_*z,i*_ and *Â*_*z,i*_ represent the pixel counts of matched instance i in slice z for the fluorescent image patch x and the in silico labeled image patch 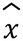, respectively.

*l*_*xy*_ = 0. 29(*µm*/*pixel*). Area (*µm*) is averaged over all matched instances *M*_*z*_ in slice z and then over all valid slices S of the 3D image patch.

(10) Hausdorff distance (“HD”) ^41^ evaluates morphological boundary and shape fidelity.

For matched instances between the fluorescent and in silico labeled images, we calculate the slice-level Hausdorff distance *d*_*H*_, defined by the maximum nearest-point distance between the sets of points representing the instances’ contours:

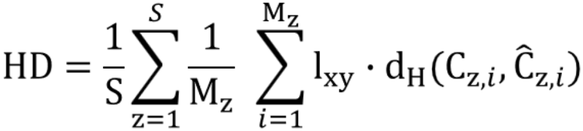

Where *C*_*z,i*_ and *Ĉ*_*z,i*_ represent the contours of the matched instance i in slice z for the fluorescent image patch x and the in silico labeled image patch 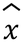, respectively.

*l*_*xy*_ = 0. 29(*µm*/*pixel*). HD (*µm*) is averaged over all matched instances *M*_*z*_ in slice z and then over all valid slices S of the 3D image patch.

### Benchmarking visualization and statistical analysis

We evaluated the relative performance of the DM and the DM4ISL over the ten metrics by calculating the ratio between the averages across all test image patches of each model to the other models. *metric_sign* was set to -1 for the MSE, Centroid distance, Neighbor distance, Area and HD metrics, for which lower values indicate better performance.

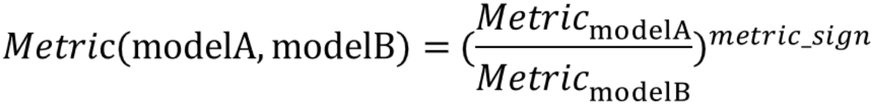

Where modelA is DM or DM4ISL and modelB can be DM, U-Net or GAN.

To calculate the overall relative performance of the DM and DM4ISL models we averaged the relative performance ratios across organelles for each metric, and then across metrics. For example, a ratio of 1.085 (8.5%) was shown between DM4ISL and DM overall metrics and all organelles.

To visualize the benchmarking of the four in silico models over these ten metrics we generated relative performance polar plots. Each metric was scaled by setting the best model to a value of 1. Specifically, we identified the highest-performing model, 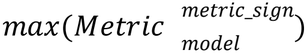.

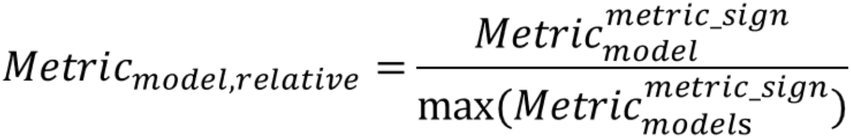

Statistical assessment was performed using the Wilcoxon signed-rank test, a non-parametric approach suitable for the non-normal distributions observed in our image similarity metrics. The null hypothesis assumed the median difference between paired metric values across the test image patches was zero. A two-sided P < 0.05 was considered statistically significant.

### Expert assessment of unambiguous substantial errors

An expert (O.R.) manually annotated all image patches in the test set that exhibited unambiguous substantial errors in the Attention U-net and in the DM4ISL in silico labeling predictions. Every z-stack slice of each in silico labeled image patch was visually compared to its respective ground-truth fluorescence. An image patch was considered as exhibiting unambiguous substantial errors if it included a visually apparent deviation in a predicted organelle’s localization, shape, or intensity. Unambiguous substantial errors in the DNA were further categorized into “deformed”, “omitted nucleus”, and “hallucinated nucleus”. Because mitochondrial structures are highly dense and complex, their predictions were excluded from this manual analysis to avoid unreliable visual assessments.

### Uncertainty estimation

We used DM4ISL’s spatial standard deviation image 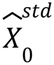, that reflects the model’s local prediction instability, to define our uncertainty estimation measurement. The standard deviation images 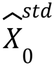 underwent a processing pipeline to result in an uncertainty map. For each organelle we defined the 99^th^ percentile over all pixels in all 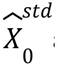 across the test set as a noise removal threshold, to remove very weak intensities. Specifically, the threshold for the Nuclear envelope was set to 0.055, for the Nucleoli to 0.065, for the Actin filaments to 0.09 and for the Microtubules to 0.04. The DNA exhibited a higher number of manually annotated errors, and thus we decided to set the uncertainty estimation threshold to the 94th percentile (0.045) to increase sensitivity. Next a median filter with kernel 3×3 was used to smooth the remaining noise and edges. Next Otsu thresholding was used to segment each image patch separately using all slice pixels. Finally, small objects, with areas of 15 pixels^2 or less, were removed and holes were filled using a 3×3 kernel at the slice-level.

This process is depicted in Fig. 4D. Finally, we defined the uncertainty estimation as the percentage of positive (high uncertainty) pixels in the uncertainty map. A threshold of uncertainty estimation in at least 1% of a patch’s pixels was set to flag potential in silico labeling errors. Mitochondria were also excluded from this uncertainty analysis.

### Prediction error annotation and uncertainty benchmark dataset

The open-access benchmark dataset comprises 128 held-out 3D test patches for each of the six organelles (DNA, Nuclear envelope, Nucleoli, Actin filaments, Mitochondria, and Microtubules) cropped from the Allen Institute hiPSC dataset. Each entry includes paired brightfield and fluorescence volumes, and corresponding predictions generated by DM, DM4ISL, U-Net, and GAN models. For five organelles (excluding mitochondria), the dataset also includes per-pixel standard-deviation maps, binary uncertainty maps, and expert manual error annotations for U-Net and DM4ISL predictions, as well as the evaluation of the DM4ISL uncertainty maps against these annotations.

### DREMI visualization

Standard mutual information fails to capture the true relationship between IoU scores and uncertainty estimates because the raw data points cluster overwhelmingly within a small, localized region (Fig. 4J). To prevent these dense clusters from dominating the visualization, we calculated the Density Resampling Estimate of Mutual Information (DREMI) ^43^. DREMI neutralizes density imbalances by discretizing the uncertainty estimate spectrum into uniform intervals and evaluating the IoU measurements within each bin independently. We used Kernel Density Estimation (KDE) to model the smoothed conditional and joint probability spaces.

DREMI = H(Y) - H(Y|X)

where *X* represents the uncertainty estimate and *Y* represents the IoU measurement, H(Y) is the marginal entropy of the IoU measurements, and H(Y|X) is the conditional entropy of the IoU given the uncertainty estimate.

### Computational resources

The source code was implemented in Python 3.9 with PyTorch 2.2 and CUDA 11.8. The models were trained on a single NVIDIA RTX 6000 GPU.

## Data and Code Availability

DM4ISL source code is publicly available via GitHub at https://github.com/zaritskylab/DM4ISL. The full data of paired brightfield and fluorescence images can be downloaded from the Allen Institute for Cell Science https://open.quiltdata.com/b/allencell/tree/aics/pipeline_integrated_cell/fovs/. Code for downloading and organizing the Allen Institute data according to the file naming and channel structure expected by our pipeline is provided in our repository. The prediction error annotation and uncertainty benchmark dataset, including paired brightfield and fluorescence test patches, model predictions, uncertainty maps, and expert prediction-error annotations is publicly available from the BioImage Archive ^67,68^ under accession S-BIAD3820 ^67,68^. Trained DM4ISL models for all six organelles, the train/test FOV lists used in the study, and example data for running the accompanying code are available on Zenodo ^69^.

## Funding and Acknowledgements

This research was supported by the Israel Science Foundation (ISF, grant no. 2516/21 to A.Z), by the Israeli Council for Higher Education (CHE) via the Data Science Research Center, Ben-Gurion University of the Negev, Israel (to A.Z.) and by the Allen Family Philanthropies (to A.Z.). We thank Noga Levy for critical reading of the manuscript. We thank Leon Ben Nedava for downloading and organizing the data from the Allen Institute for Cell Science.

## Author Contribution Statement

A.Z. and O.R. conceived the study. O.R. developed the computational method and analyzed the data. O.K.G. revised and validated the computational analyses and organized the code and data for sharing. O.R. and A.Z. interpreted the data and drafted the manuscript. A.Z. mentored O.R. All authors edited the manuscript and approved its content.

## Competing Interests Statement

The authors declare no competing interests.

## Supplementary Information

### Supplementary Figures

**Figure S1.**
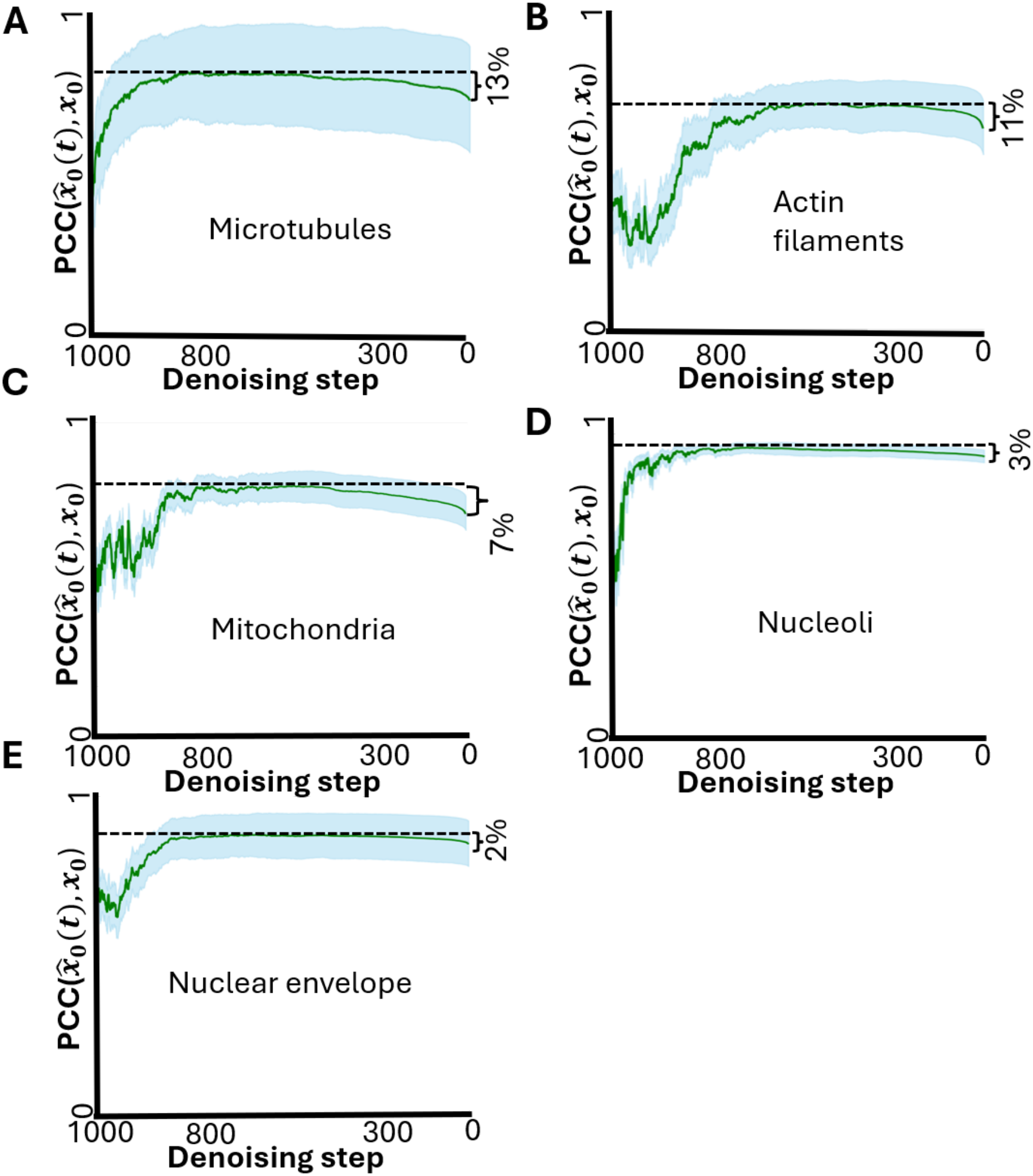
Deteriorated performance in the late stages of denoising. The evolution of the mean (green) and the standard deviation (cyan) PCC between the fluorescence prediction 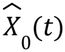 and the corresponding fluorescence images *X*_0_ throughout the iterative denoising process. The horizontal dashed black line depicts the optimal 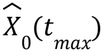 emphasizing a deterioration of the DM’s final fluorescence prediction 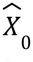. **(A)** Microtubules, 13% deterioration. **(B)** Actin filaments, 11% deterioration. **(C)** Mitochondria, 7% deterioration. **(D)** Nucleoli, 3% deterioration. **(E)** Nuclear envelope, 2% deterioration.

**Figure S2.**
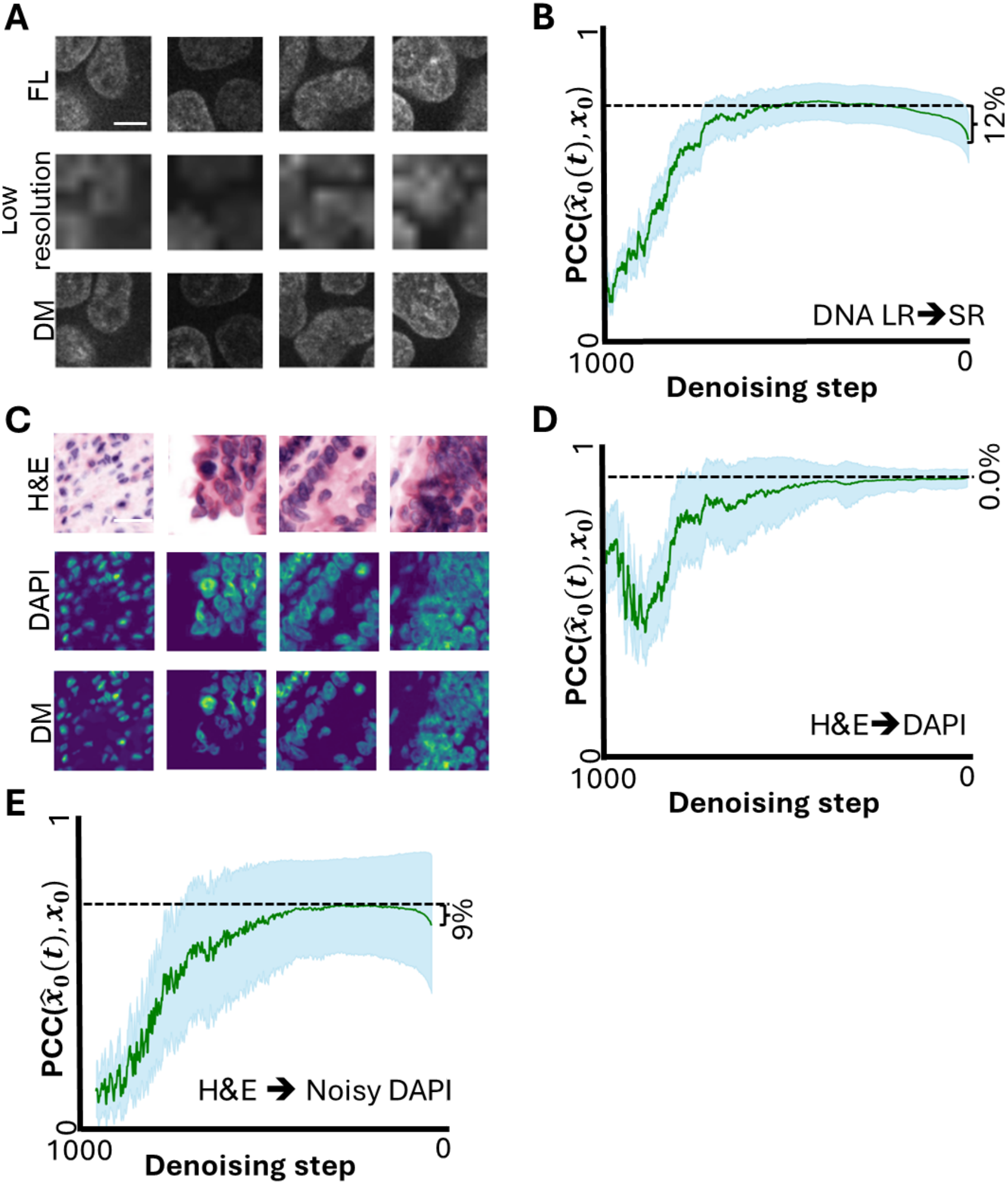
Super-resolution fluorescence reconstruction and H&E-to-DAPI translation. **(A)** Super-resolution of four image patches. Top-to-bottom: the DNA fluorescence image patches, downsampling to reduce resolution (Methods), the DM’s 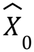 prediction. Scale bar = 10μm. **(B)** The evolution of the mean (green) and the standard deviation (cyan) Pearson correlation coefficient (PCC) between the DNA’s super-resolution fluorescence prediction 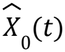 and the corresponding fluorescence images *X*_0_ throughout the iterative denoising process. The horizontal dashed black line depicts the optimal 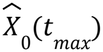 emphasizing a deterioration of 12% in the DM’s final fluorescence prediction 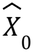. **(C)** H&E-to-DAPI translation of four image patches. Top-to-bottom: the H&E image patches, the corresponding DAPI stained images, the DM’s 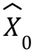. Scale bar = 100μm. **(D)** The evolution of the mean (green) and the standard deviation (cyan) Pearson correlation coefficient (PCC) between the DAPI prediction 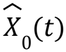 and the corresponding DAPI images *X* throughout the iterative denoising process. The horizontal dashed black line depicts the optimal 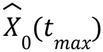occurs for 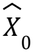, emphasizing no deterioration. **(E)** Same as D, after inclusion of Gaussian and Poisson noise to the DAPI images (Methods).

**Figure S3.**
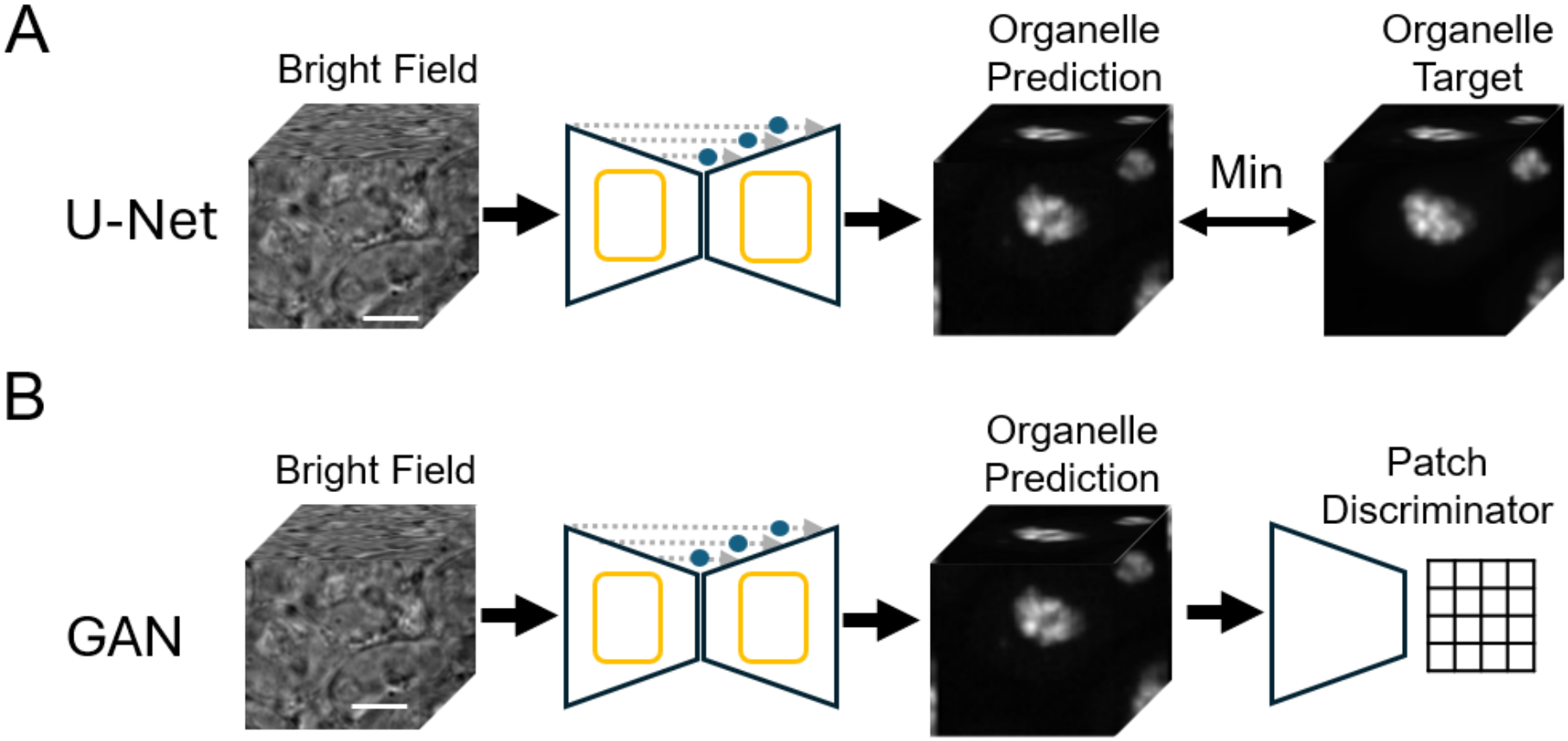
Common architectures for cross modality image translation. **(A)** Attention U-Net. The brightfield image patches are processed through an Attention U-Net architecture, where integrated attention gates (blue circles) selectively highlight salient feature maps to predict the corresponding fluorescence patches. The model is trained by minimizing the mean absolute error (MAE) loss between the in silico predictions and the ground-truth fluorescence patches. **(B)** Pix2pix GAN. The Attention U-Net architecture backbone is employed as the generator network within the Pix2Pix framework. A patch discriminator is used to distinguish ground-truth fluorescence from generator predictions by outputting a 4×4×4 grid of probabilities, where each cell represents the likelihood that a specific image partition is authentic. The model is optimized using a combination of the generator’s pixel-wise MAE loss and an adversarial loss derived from the discriminator’s ability to classify real versus synthetic predictions. Scale bars = 10μm.

**Figure S4.**
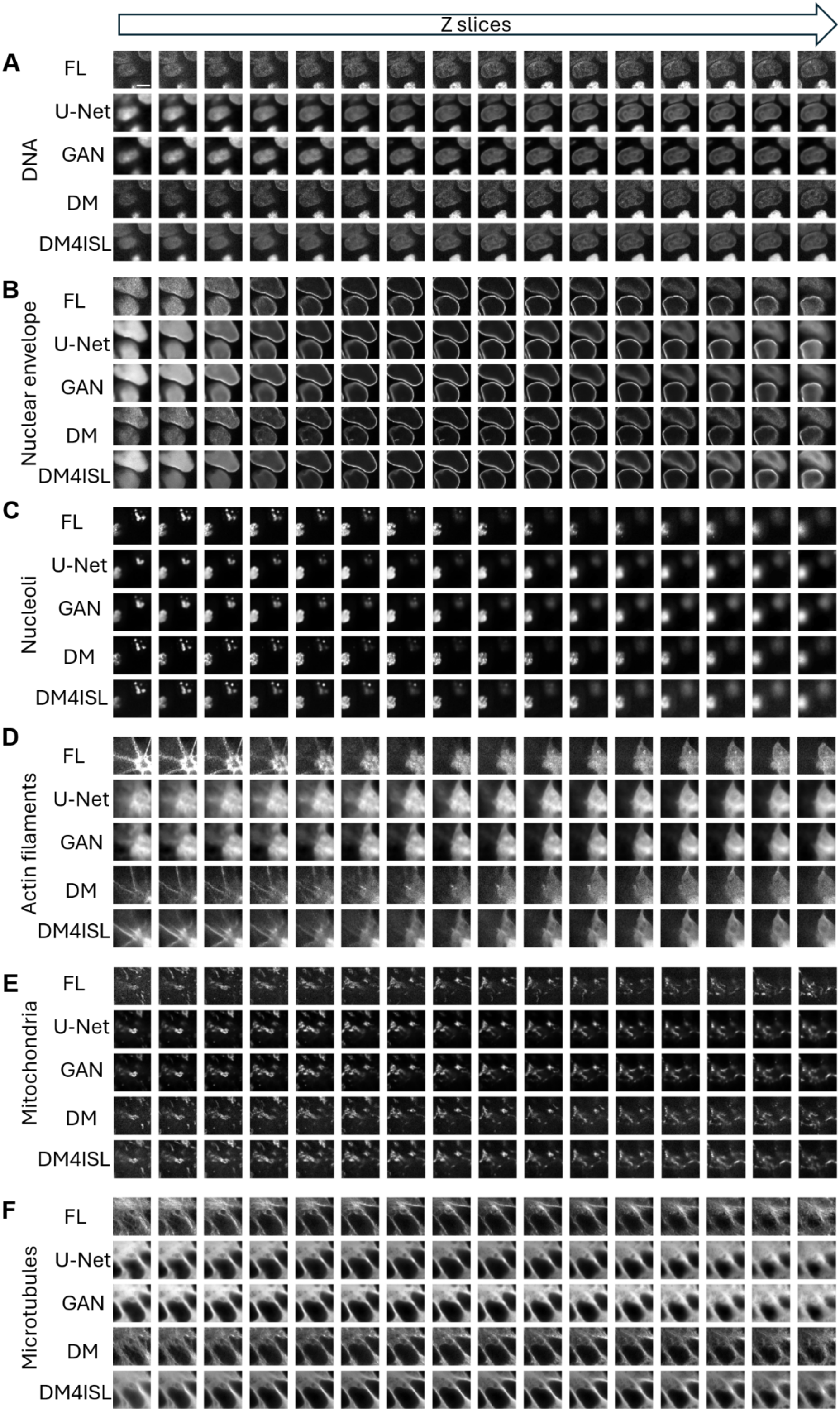
Visual qualitative assessment of different in silico labeling models in 3D. Axial slices from a representative 3D field of view for each organelle (columns). Rows display (top-to-bottom): the ground truth fluorescence image, the corresponding attention U-Net, Pix2Pix GAN, DM and DM4ISL models. (**A**) DNA. (**B**) Nuclear envelope. (**C**) Nucleoli. (**D**) Actin filaments. (**E**) Mitochondria. (**F**) Microtubules. Scale bars = 10μm.

**Figure S5.**
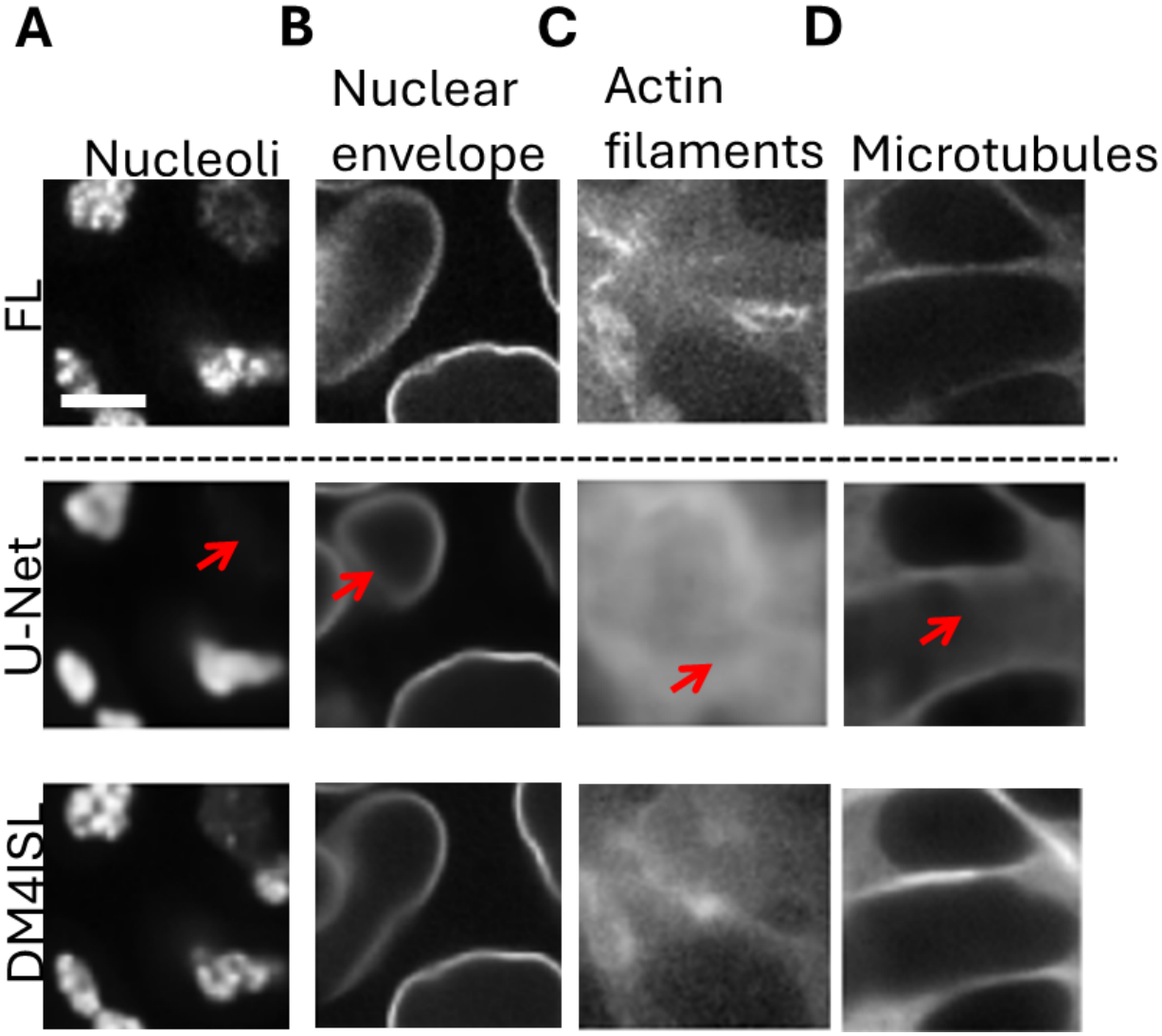
Representative erroneous predictions of the Nucleoli, Nuclear envelope, Actin filaments and Microtubules by the U-Net. Top-to-bottom: fluorescence image patches (‘FL’); the corresponding Attention U-Net’s (‘U-Net’) in silico labeling; and the corresponding DM4ISL’s in silico labeling. Red arrows guide to regions where the U-Net, substantially deviates from the fluorescence image (columns). **(A)** Nucleoli. **(B)** Nuclear envelope. **(C)** Actin filaments. **(D)** Microtubules. Errors in Mitochondria prediction were not annotated because we were not able to conclusively annotate what were substantial unambiguous errors. Scale bar = 10μm.

**Figure S6.**
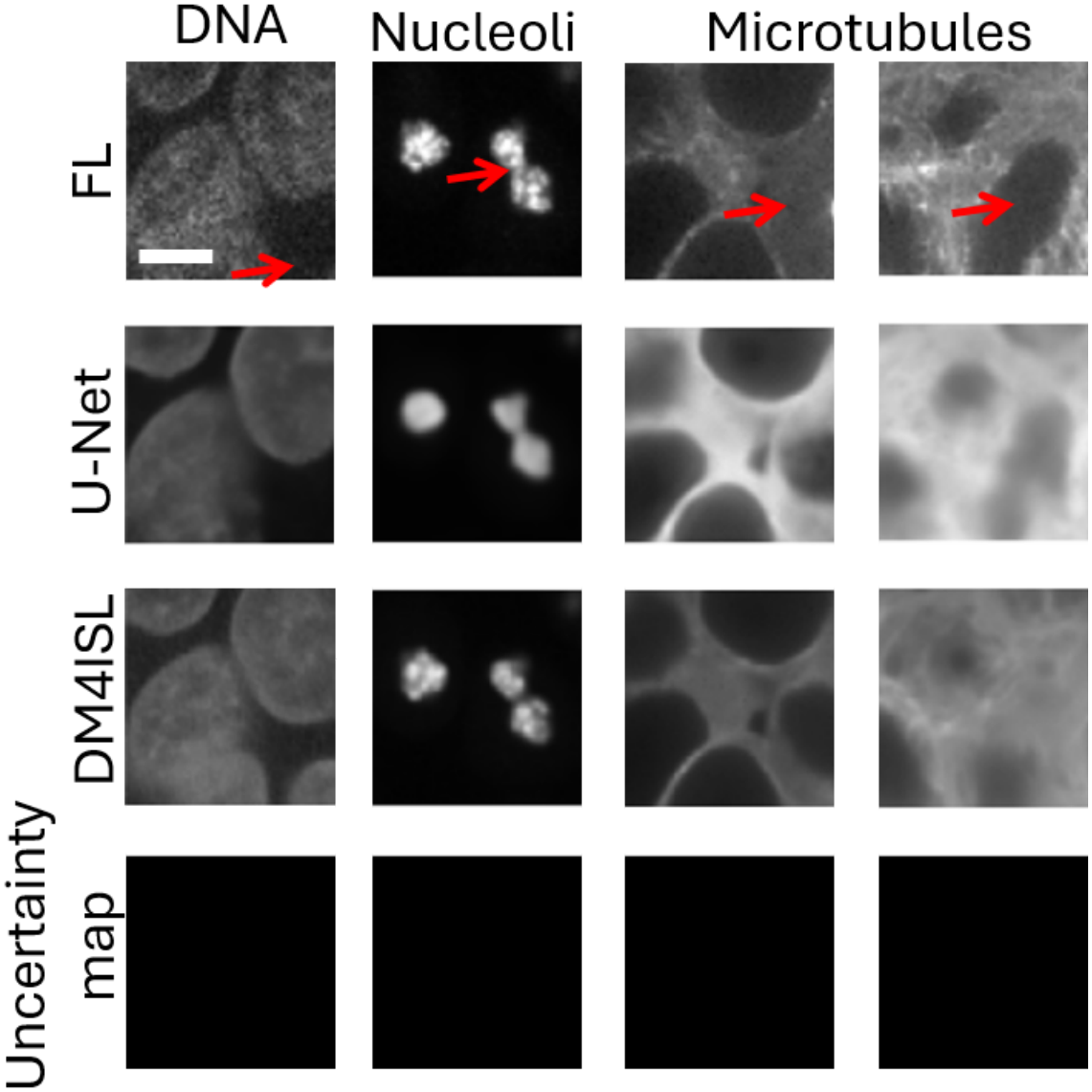
Substantial errors missed by the DM4ISL-derived uncertainty maps. DM4ISL-derived uncertainty maps miss 4 manually annotated substantial errors across organelle (columns): one for the DNA, one for the Nucleoli, and two for the Microtubules. Row (top-to-bottom): the fluorescence image patch (FL), the corresponding U-Net fluorescence prediction (U-Net), the corresponding DM4ISL fluorescence prediction (DM4ISL), and the corresponding uncertainty map. Red arrows guide to regions where the uncertainty map missed a manually annotated substantial error. Scale bar = 10μm.

**Figure S7.**
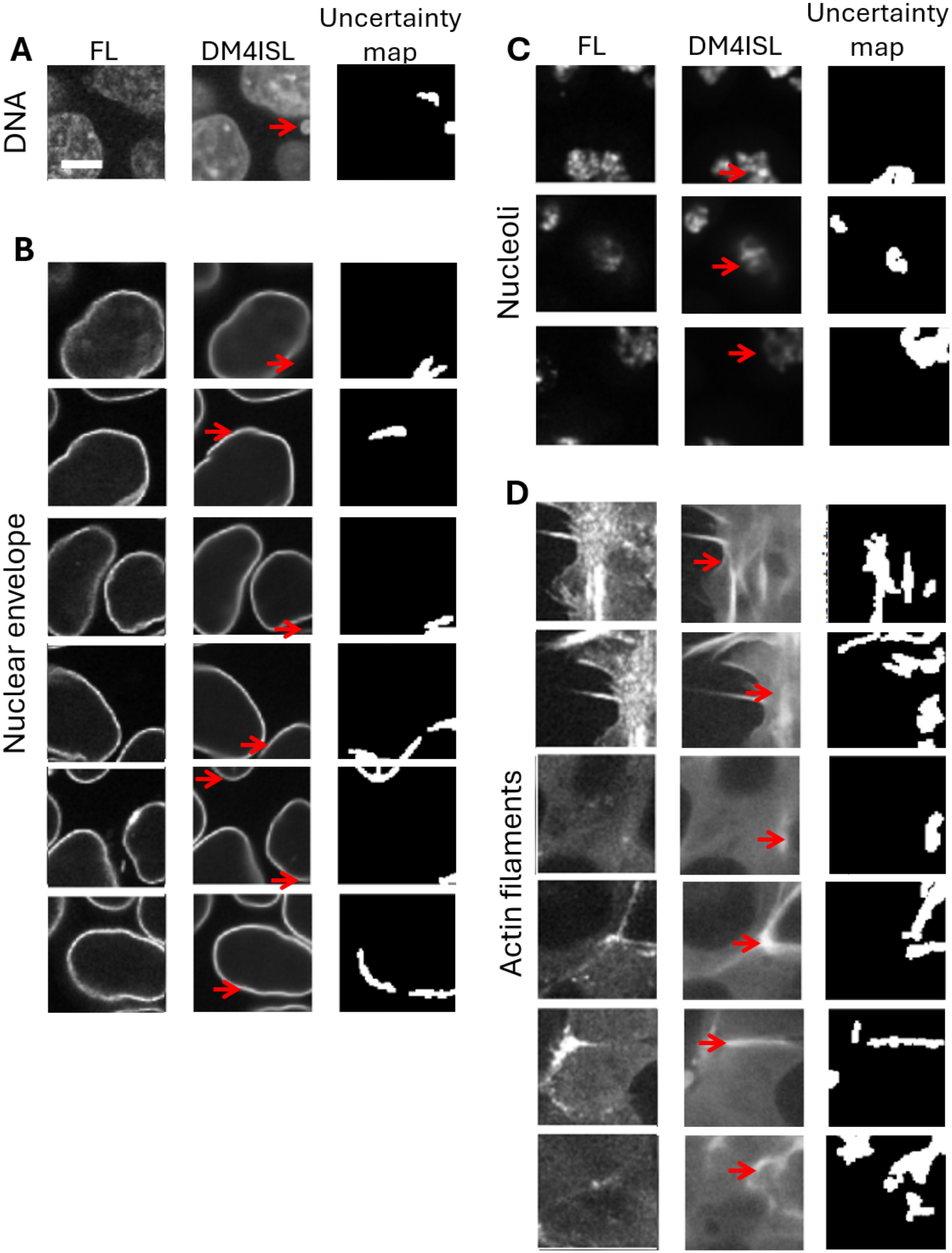
DM4ISL-derived uncertainty maps flagged in silico labeling errors that were missed during our manual annotation. Each panel represents an organelle, displaying image patches that were correctly flagged by the corresponding uncertainty maps after being overlooked during the manual inspection. Left-to-right: the fluorescence image patch (FL), the corresponding DM4ISL fluorescence prediction (DM4ISL), and the corresponding uncertainty map. **(A)** DNA. **(B)** Nuclear envelope. **(C)** Nucleoli. **(D)** Actin filaments. Red arrows guide to regions where the DM4ISL’s prediction substantially deviates from the fluorescence image. Scale bar = 10μm.

**Figure S8.**
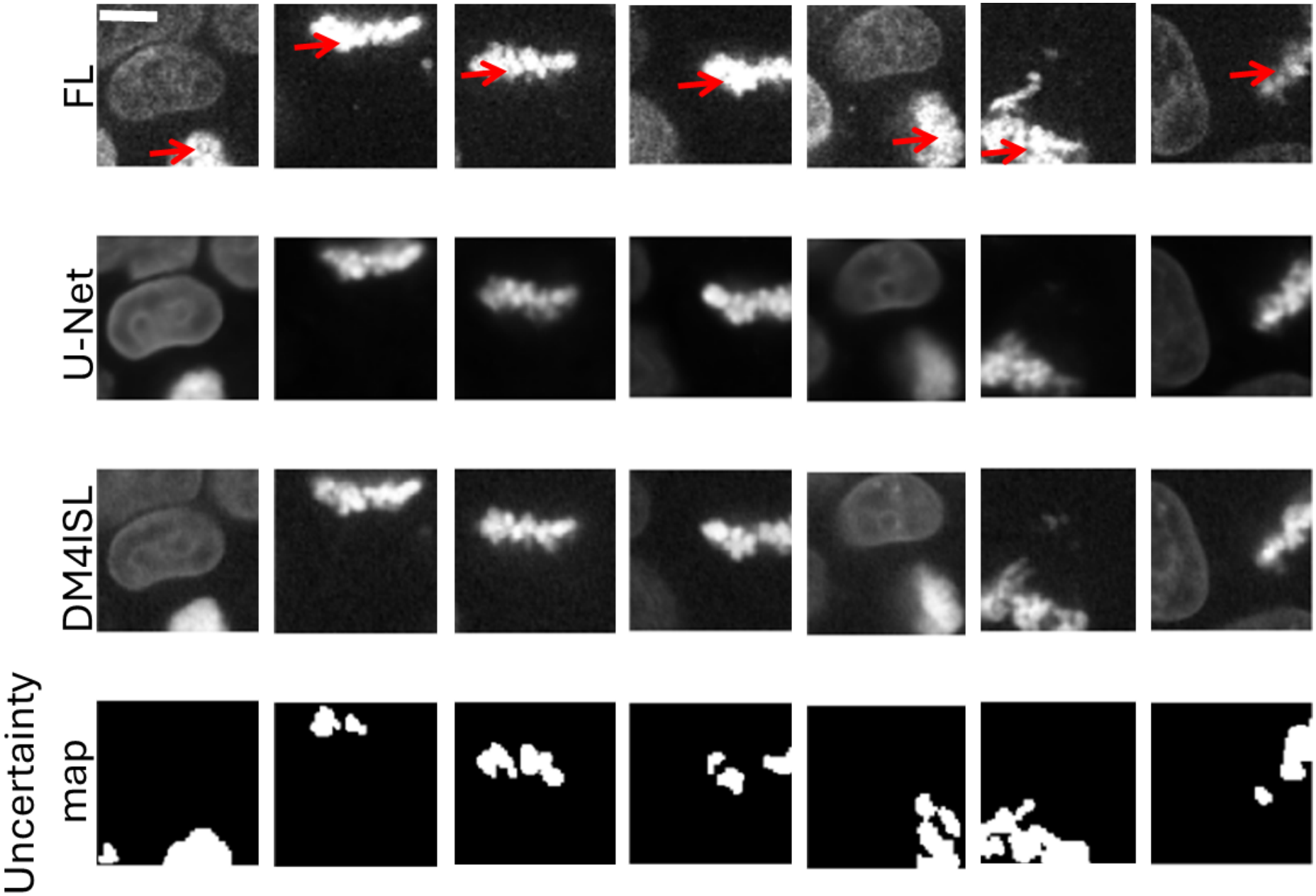
DM4ISL-derived uncertainty estimated DNA false alarms flag abnormal cell states. Representative DNA image patches that were not manually annotated as prediction errors, but were flagged by the DM4ISL-derived uncertainty estimates. Most of these false alarms were attributed to cells with abnormal shape that were also associated with deteriorated application-appropriate in silico labeling (Fig. 4J). Top-to-bottom: the fluorescence image patch (FL), the corresponding U-Net fluorescence prediction (U-Net), the corresponding DM4ISL fluorescence prediction (DM4ISL), and the corresponding uncertainty maps. Red arrows guide to regions of the abnormal regions in the DNA fluorescence image. Scale bar = 10μm.

### Supplementary Tables

**Table S1.**
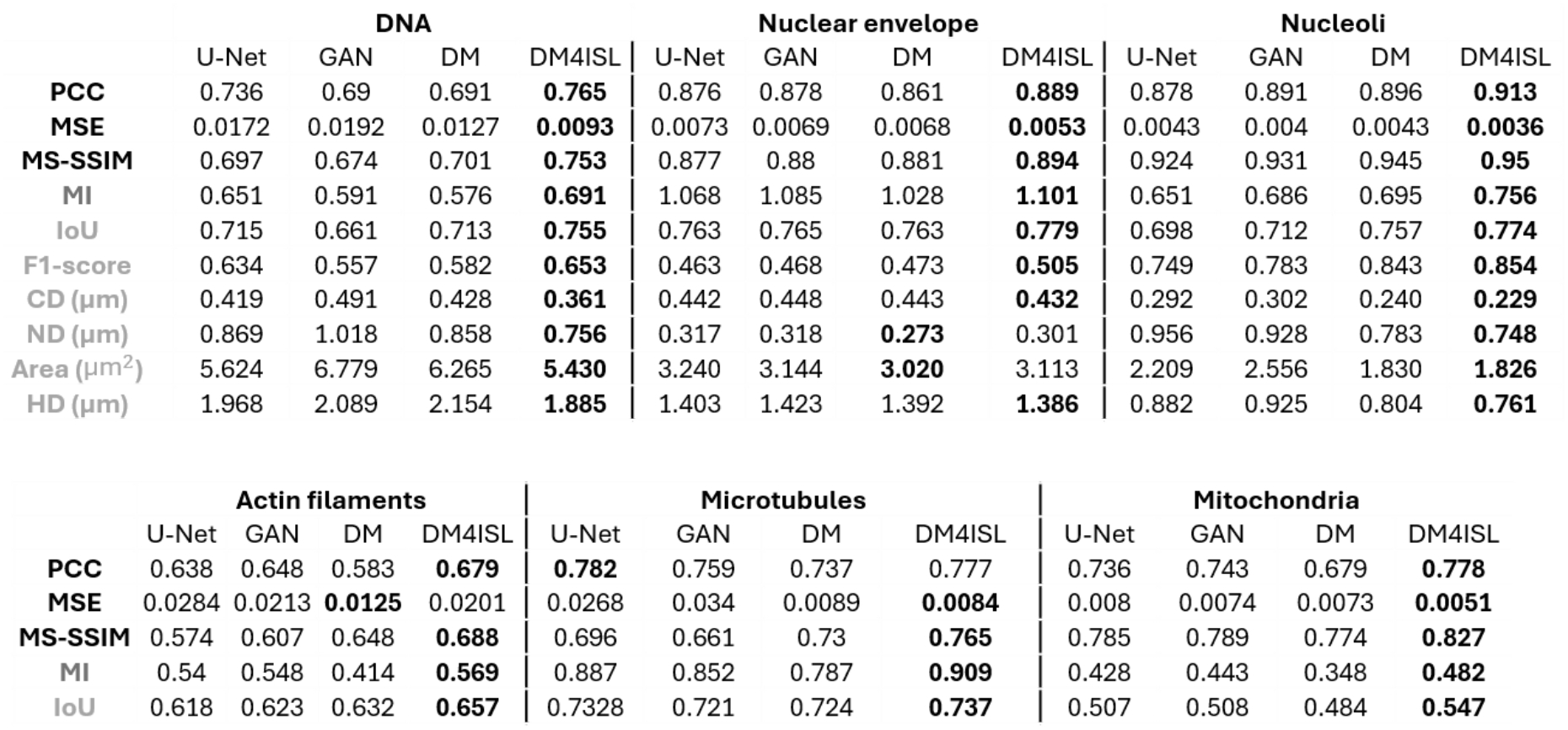
**Benchmarking**. Four in silico labeling models were benchmarked across six organelles using ten metrics. Each organelle was evaluated according to n = 128 image patches randomly selected from 8-19 fields of view. Three types of image quality evaluation metrics: pixel and localized patch-scales (black), intensity distribution (dark gray), and application-appropriate measurements (light gray). Higher values indicate superior performance for the metric MS-SSIM, PCC, MI, IoU and F1, and lower values indicate superior performance for MSE, CD, ND, Area and HD. The best performing model for each organelle-metric pair is marked in bold. DM4ISL was the best performing model for 41 of the 45 (∼91%) organelle-metric pairs.

|  | **DNA** |  |  |  | **Nuclear envelope** |  |  |  | **Nucleoli** |  |  |  |
|---|---|---|---|---|---|---|---|---|---|---|---|---|
|  | U-Net | GAN | DM | DM4ISL | U-Net | GAN | DM | DM4ISL | U-Net | GAN | DM | DM4ISL |
| **PCC** | 0.736 | 0.69 | 0.691 | **0.765** | 0.876 | 0.878 | 0.861 | **0.889** | 0.878 | 0.891 | 0.896 | **0.913** |
| **MSE** | 0.0172 | 0.0192 | 0.0127 | **0.0093** | 0.0073 | 0.0069 | 0.0068 | **0.0053** | 0.0043 | 0.004 | 0.0043 | **0.0036** |
| **MS-SSIM** | 0.697 | 0.674 | 0.701 | **0.753** | 0.877 | 0.88 | 0.881 | **0.894** | 0.924 | 0.931 | 0.945 | **0.95** |
| **MI** | 0.651 | 0.591 | 0.576 | **0.691** | 1.068 | 1.085 | 1.028 | **1.101** | 0.651 | 0.686 | 0.695 | **0.756** |
| **IoU** | 0.715 | 0.661 | 0.713 | **0.755** | 0.763 | 0.765 | 0.763 | **0.779** | 0.698 | 0.712 | 0.757 | **0.774** |
| **F1-score** | 0.634 | 0.557 | 0.582 | **0.653** | 0.463 | 0.468 | 0.473 | **0.505** | 0.749 | 0.783 | 0.843 | **0.854** |
| **CD (µm)** | 0.419 | 0.491 | 0.428 | **0.361** | 0.442 | 0.448 | 0.443 | **0.432** | 0.292 | 0.302 | 0.240 | **0.229** |
| **ND (µm)** | 0.869 | 1.018 | 0.858 | **0.756** | 0.317 | 0.318 | **0.273** | 0.301 | 0.956 | 0.928 | 0.783 | **0.748** |
| **Area** (µm<sup>2</sup>) | 5.624 | 6.779 | 6.265 | **5.430** | 3.240 | 3.144 | **3.020** | 3.113 | 2.209 | 2.556 | 1.830 | **1.826** |
| **HD (µm)** | 1.968 | 2.089 | 2.154 | **1.885** | 1.403 | 1.423 | 1.392 | **1.386** | 0.882 | 0.925 | 0.804 | **0.761** |
|  | **Actin filaments** |  |  |  | **Microtubules** |  |  |  | **Mitochondria** |  |  |  |
|  | U-Net | GAN | DM | DM4ISL | U-Net | GAN | DM | DM4ISL | U-Net | GAN | DM | DM4ISL |
| **PCC** | 0.638 | 0.648 | 0.583 | **0.679** | **0.782** | 0.759 | 0.737 | 0.777 | 0.736 | 0.743 | 0.679 | **0.778** |
| **MSE** | 0.0284 | 0.0213 | **0.0125** | 0.0201 | 0.0268 | 0.034 | 0.0089 | **0.0084** | 0.008 | 0.0074 | 0.0073 | **0.0051** |
| **MS-SSIM** | 0.574 | 0.607 | 0.648 | **0.688** | 0.696 | 0.661 | 0.73 | **0.765** | 0.785 | 0.789 | 0.774 | **0.827** |
| **MI** | 0.54 | 0.548 | 0.414 | **0.569** | 0.887 | 0.852 | 0.787 | **0.909** | 0.428 | 0.443 | 0.348 | **0.482** |
| **IoU** | 0.618 | 0.623 | 0.632 | **0.657** | 0.7328 | 0.721 | 0.724 | **0.737** | 0.507 | 0.508 | 0.484 | **0.547** |

**Table S2.**
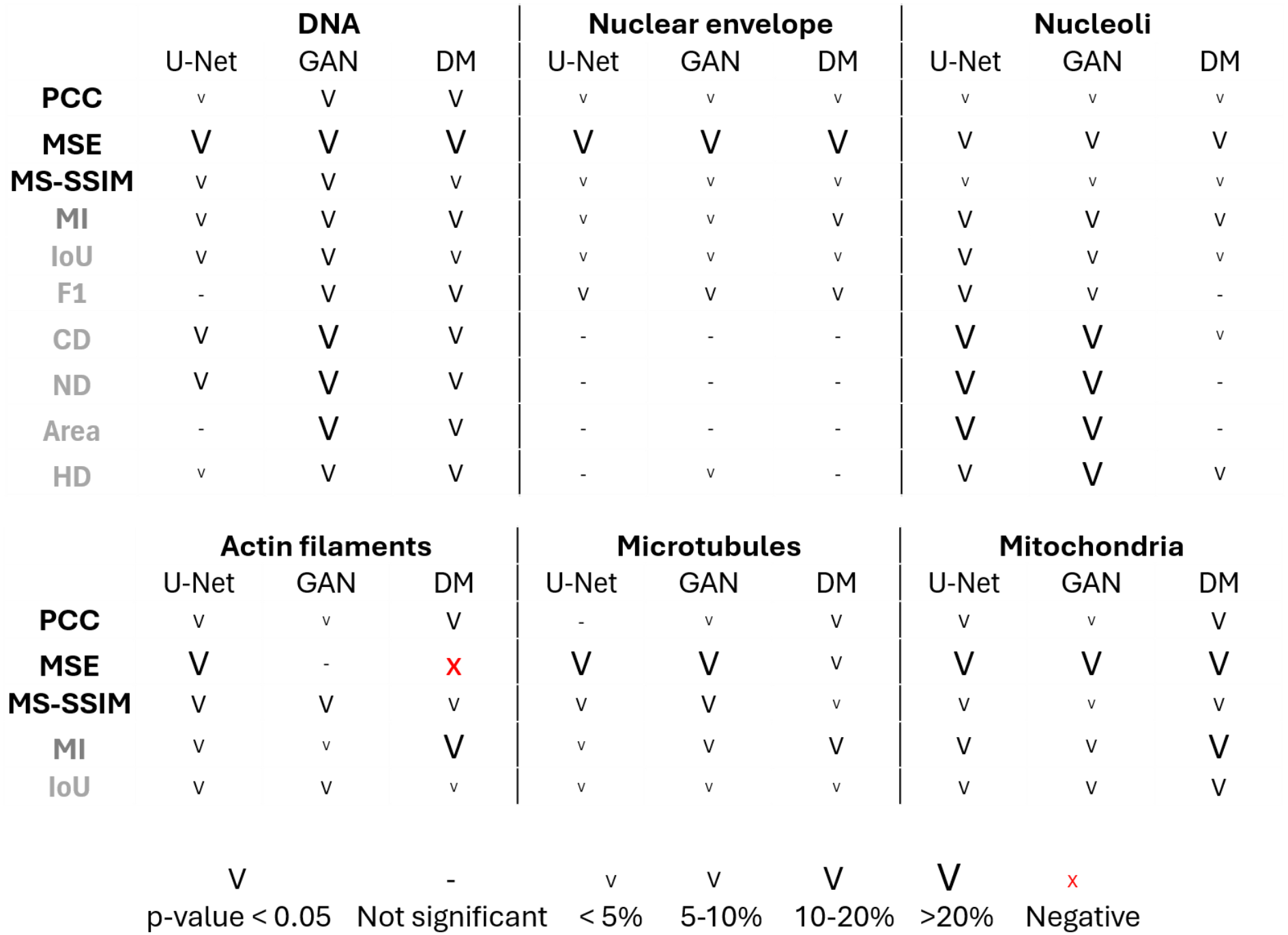
Summary of DM4ISL performance gain and statistical significance versus the alternative models. Each bin reports the performance gain of DM4ISL and the corresponding statistical significance for a given organelle (panel) and metric (row), compared to an alternative model (columns), according to the benchmarking in Table S1. The marker ‘V’ indicates statistical significance (p-value < 0.05) according to the two-sided Wilcoxon signed-rank test. Font size reflects the magnitude of DM4ISL gain. The marker ‘X’ indicates statistically significant diminished performance of DM4ISL relative to the competing model. The marker ‘-’ indicates non-significance.

|  | DNA |  |  | Nuclear envelope |  |  | Nucleoli |  |  |
|---|---|---|---|---|---|---|---|---|---|
|  | U-Net | GAN | DM | U-Net | GAN | DM | U-Net | GAN | DM |
| **PCC** | v | V | V | v | v | v | v | v | v |
| **MSE** | V | V | V | V | V | V | V | V | V |
| **MS-SSIM** | V | V | V | v | v | v | v | v | v |
| **MI** | V | V | V | v | v | V | V | V | V |
| **IoU** | V | V | V | v | v | v | V | V | v |
| **F1** | - | V | V | V | V | V | V | V | - |
| **CD** | V | V | V | - | - | - | V | V | v |
| **ND** | V | V | V | - | - | - | V | V | - |
| **Area** | - | V | V | - | - | - | V | V | - |
| **HD** | v | V | V | - | v | - | V | V | V |
|  | **Actin filaments** |  |  | **Microtubules** |  |  | **Mitochondria** |  |  |
|  | U-Net | GAN | DM | U-Net | GAN | DM | U-Net | GAN | DM |
| **PCC** | V | v | V | - | v | V | V | v | V |
| **MSE** | V | - | x | V | V | V | V | V | V |
| **MS-SSIM** | V | V | V | V | V | v | V | v | V |
| **MI** | V | v | V | v | V | V | V | V | V |
| **IoU** | V | V | v | v | v | v | V | V | V |

V p-value < 0.05 - Not significant v < 5% V 5-10% V 10-20% V >20% x Negative

**Table S3.** Robustness of the interval of the range of intermediate denoising steps. Sensitivity analysis for the range of DM4ISL’s intermediate denoising steps performed on a subset of the Nucleoli test set.

| **Nucleoli** |  |  |
|---|---|---|
| t low | t high | PCC |
| 200 | 700 | 0.8762 |
| 200 | 800 | 0.8768 |
| 250 | 800 | 0.8766 |
| 300 | 800 | 0.8763 |
| 350 | 800 | 0.8760 |

## Notes

https://www.ebi.ac.uk/biostudies/bioimages/studies/S-BIAD3820

https://zenodo.org/records/21921959

https://github.com/zaritskylab/DM4ISL

### Competing Interest Statement

The authors have declared no competing interest.

## References

1. Sullivan, K. F. & Kay, S. A. (eds) Green Fluorescent Proteins. Methods in Cell Biology Vol. 58 (Academic Press, 1999).

2. Garini, Y., Young, I. T. & McNamara, G. Spectral imaging: principles and applications. Cytometry A 69, 735–747 (2006).

3. Icha, J., Weber, M., Waters, J. C. & Norden, C. Phototoxicity in live fluorescence microscopy, and how to avoid it. Bioessays 39, 1700003 (2017).

4. Scherf, N. & Huisken, J. The smart and gentle microscope. Nat Biotechnol 33, 815–818 (2015).

5. Elmalam, N., Ben Nedava, L. & Zaritsky, A. In silico labeling in cell biology: Potential and limitations. Curr Opin Cell Biol 89, 102378 (2024).

6. Ounkomol, C., Seshamani, S., Maleckar, M. M., Collman, F. & Johnson, G. R. Label-free prediction of three-dimensional fluorescence images from transmitted-light microscopy. Nat Methods 15, 917–920 (2018).

7. Christiansen, E. M. et al. In Silico Labeling: Predicting Fluorescent Labels in Unlabeled Images. Cell 173, 792–803.e19 (2018).

8. Jo, Y. et al. Label-free multiplexed microtomography of endogenous subcellular dynamics using generalizable deep learning. Nat Cell Biol 23, 1329–1337 (2021).

9. Cheng, S. et al. Single-cell cytometry via multiplexed fluorescence prediction by label-free reflectance microscopy. Sci Adv 7, eabe0431 (2021).

10. Wang, Z., Xie, Y. & Ji, S. Global voxel transformer networks for augmented microscopy. Nature Machine Intelligence 3, 161–171 (2021).

11. Liu, Z. et al. Robust virtual staining of landmark organelles with Cytoland. Nature Machine Intelligence 7, 901–915 (2025).

12. Follain, G. et al. Fast label-free live imaging with FlowVision reveals key principles of cancer cell arrest on endothelial monolayers. EMBO J. 45, 1381–1421 (2026).

13. Kalinin, A. A. et al. Foreground-aware Virtual Staining for Accurate 3D Cell Morphological Profiling. Preprint at arXiv 10.48550/arXiv.2507.05383 (2025).

14. Nichol, A. & Dhariwal, P. Improved Denoising Diffusion Probabilistic Models. In Proc. 38th International Conference on Machine Learning Vol. 139, 8162–8171 (PMLR, 2021).

15. Ho, J., Jain, A. & Abbeel, P. Denoising Diffusion Probabilistic Models. In Advances in Neural Information Processing Systems 33 6840–6851 (Curran Associates, 2020).

16. Saharia, C. et al. Palette: Image-to-Image Diffusion Models. In ACM SIGGRAPH 2022 Conference Proceedings 15 (ACM, 2022).

17. Wolleb, J., Bieder, F., Sandkühler, R. & Cattin, P. C. Diffusion Models for Medical Anomaly Detection. In Medical Image Computing and Computer Assisted Intervention – MICCAI 2022 Lecture Notes in Computer Science Vol. 13438, 35–45 (Springer, 2022).

18. Wu, J. et al. MedSegDiff: Medical Image Segmentation with Diffusion Probabilistic Model. In Proc. Medical Imaging with Deep Learning Vol. 227 (PMLR, 2024).

19. Cahan, N. et al. X-ray2CTPA: leveraging diffusion models to enhance pulmonary embolism classification. NPJ Digit Med 8, 439 (2025).

20. Li, R. et al. Microscopy image reconstruction with physics-informed denoising diffusion probabilistic model. Communications engineering 3, 186 (2024).

21. Yan, J., Tang, Z., Yan, Q. & Huang, S. Prior-enhanced diffusion model for super-resolution reconstruction in structured illumination microscopy. Optics and Lasers in Engineering 197, 109468 (2026).

22. Saguy, A. et al. This Microtubule Does Not Exist: Super-Resolution Microscopy Image Generation by a Diffusion Model. Small Methods 9, e2400672 (2025).

23. Acuña-Rodriguez, J. P., Mena-Vega, J. P. & Argüello-Miranda, O. Live-cell fluorescence spectral imaging as a data science challenge. Biophys. Rev. 14, 579–597 (2022).

24. Fazel, M. et al. Fluorescence Microscopy: a statistics-optics perspective. Rev. Mod. Phys. 96, 025003 (2024).

25. Chen, J., Viana, M. P. & Rafelski, S. M. When seeing is not believing: application-appropriate validation matters for quantitative bioimage analysis. Nat Methods 20, 968–970 (2023).

26. Viana, M. P. et al. Integrated intracellular organization and its variations in human iPS cells. Nature 613, 345–354 (2023).

27. Sun, H., Fu, X., Abraham, S., Jin, S. & Murphy, R. F. Improving and evaluating deep learning models of cellular organization. Bioinformatics 38, 5299–5306 (2022).

28. Elmalam, N. & Zaritsky, A. Cell context-dependent in silico organelle localization in label-free microscopy images. Nat. Methods 23, 405–416 (2026).

29. Pinaya, W. H. L. et al. Generative AI for Medical Imaging: extending the MONAI Framework. Preprint at arXiv 10.48550/arXiv.2307.15208 (2023).

30. Oktay, O. et al. Attention U-Net: Learning Where to Look for the Pancreas. Preprint at arXiv 10.48550/arXiv.1804.03999 (2018).

31. Song, J., Meng, C. & Ermon, S. Denoising Diffusion Implicit Models. In International Conference on Learning Representations (ICLR, 2021).

32. Qiao, C. et al. A neural network for long-term super-resolution imaging of live cells with reliable confidence quantification. Nat Biotechnol 10.1038/s41587-025-02553-8 (2025).

33. Bian, C., Philips, B., Cootes, T. & Fergie, M. HEMIT: H&E to Multiplex-immunohistochemistry Image Translation with Dual-Branch Pix2pix Generator. In Medical Image Computing and Computer Assisted Intervention – MICCAI 2024 Workshops (eds Schroder, A. et al.) 184–197 (Springer, 2025).

34. Isola, P., Zhu, J.-Y., Zhou, T. & Efros, A. A. Image-to-Image Translation with Conditional Adversarial Networks. In Proc. IEEE Conference on Computer Vision and Pattern Recognition 1125–1134 (IEEE, 2017).

35. Terpstra, H. M. et al. PunctaFinder: An algorithm for automated spot detection in fluorescence microscopy images. Mol Biol Cell 35, mr9 (2024).

36. Wang, Z., Simoncelli, E. P. & Bovik, A. C. Multiscale structural similarity for image quality assessment. In Proc. Thirty-Seventh Asilomar Conference on Signals, Systems & Computers Vol. 2, 1398–1402 (IEEE, 2003).

37. Pluim, J. P. W., Maintz, J. B. A. & Viergever, M. A. Mutual-information-based registration of medical images: a survey. IEEE Trans Med Imaging 22, 986–1004 (2003).

38. Sengupta, S. et al. On the Utility of Virtual Staining for Downstream Applications as it relates to Task Network Capacity. Biomed. Opt. Express 16, 4224–4240 (2025).

39. Gunawan, I. et al. Image quality metrics fail to accurately represent biological information in fluorescence microscopy. Preprint at bioRxiv 2025.08.05.668508 (2025) doi:10.1101/2025.08.05.668508.

40. Vincent, L. & Soille, P. Watersheds in digital spaces: an efficient algorithm based on immersion simulations. IEEE Trans. Pattern Anal. Mach. Intell. 13, 583–598 (1991).

41. Huttenlocher, D. P., Klanderman, G. A. & Rucklidge, W. J. Comparing images using the Hausdorff distance. IEEE Trans. Pattern Anal. Mach. Intell. 15, 850–863 (1993).

42. Xing, X. et al. Can generative AI replace immunofluorescent staining processes? A comparison study of synthetically generated cellpainting images from brightfield. Comput. Biol. Med. 182, 109102 (2024).

43. Krishnaswamy, S. et al. Conditional density-based analysis of T cell signaling in single-cell data. Science 346, 1250689 (2014).

44. Meijering, E., Carpenter, A. E., Peng, H., Hamprecht, F. A. & Olivo-Marin, J. C. Imagining the future of bioimage analysis. Nature biotechnology 34, 1250–1255 (2016).

45. Eliceiri, K. W. et al. Biological imaging software tools. Nature methods 9, 697–710 (2012).

46. Moen, E. et al. Deep learning for cellular image analysis. Nature methods 16, 1233–1246 (2019).

47. Laine, R. F., Arganda-Carreras, I., Henriques, R. & Jacquemet, G. Avoiding a replication crisis in deep-learning-based bioimage analysis. Nature methods 18, 1136–1144 (2021).

48. Belthangady, C. & Royer, L. A. Applications, promises, and pitfalls of deep learning for fluorescence image reconstruction. Nature methods 16, 1215–1225 (2019).

49. Bourou, A., Castaño Segade, B., Boyer, T., Mezger, V. & Genovesio, A. Revealing Subtle Phenotypes in Small Microscopy Datasets Using Latent Diffusion Models. Preprint at arXiv 10.48550/arXiv.2502.09665 (2025).

50. Zheng, D. & Huang, B. Bridging Protein Sequences and Microscopy Images with Unified Diffusion Models. In Proc. 42nd International Conference on Machine Learning Vol. 267 (PMLR, 2025).

51. Han, Y., Jiang, J. & Pei, Y. DiffStain: Conditioned diffusion-based semantic virtual staining with mask guidance. In Medical Image Computing and Computer Assisted Intervention – MICCAI 2025 (eds Gee, J. C. et al.) 139–148 (Springer, 2025).

52. Wu, D., Zhang, Y., Yeung-Levy, S., Lundberg, E. & Fox, E. B. Uncertainty quantification for distribution-to-distribution flow matching in scientific imaging. Preprint at arXiv 10.48550/arXiv.2603.21717 (2026).

53. Blau, W. et al. Protecting scientific integrity in an age of generative AI. Proc. Natl. Acad. Sci. U. S. A. 121, e2407886121 (2024).

54. Huang, Y. et al. On the trustworthiness of generative Foundation Models: Guideline, assessment, and perspective. Preprint at arXiv 10.48550/arXiv.2502.14296 (2025).

55. Ben Nedava, L. et al. Trustworthy in silico labeling via semantic visual interpretability of image-to-image translation. Preprint at bioRxiv 10.64898/2026.08.13.744455 (2026).

56. Lakshminarayanan, B., Pritzel, A. & Blundell, C. Simple and scalable predictive uncertainty estimation using deep ensembles. In Advances in Neural Information Processing Systems Vol. 30 (eds Guyon, I. et al.) 6402–6413 (Curran Associates, 2017).

57. Imboden, S., Liu, X., Payne, M. C.Hsieh, C.-J. & Lin, N. Y. C. Trustworthy in silico cell labeling via ensemble-based image translation. Biophys Rep (N Y) 3, 100133 (2023).

58. Ray, A., Galinova, V. & Jug, F. ResMatching: Noise-Resilient Computational Super-Resolution via Guided Conditional Flow Matching. In Proc. 2026 IEEE 23rd International Symposium on Biomedical Imaging (ISBI) 1–5 (IEEE, 2026).

59. Osuna-Vargas, P. et al. Denoising diffusion models for high-resolution microscopy image restoration. In Proc. IEEE/CVF Winter Conference on Applications of Computer Vision (WACV) 4320–4330 (2025).

60. Lu, C. et al. Diffusion-based deep learning method for augmenting ultrastructural imaging and volume electron microscopy. Nat. Commun. 15, 4677 (2024).

61. Rombach, R., Blattmann, A., Lorenz, D., Esser, P. & Ommer, B. High-Resolution Image Synthesis with Latent Diffusion Models. In Proc. IEEE/CVF Conference on Computer Vision and Pattern Recognition 10684–10695 (IEEE, 2022).

62. Esser, P. et al. Scaling Rectified Flow Transformers for High-Resolution Image Synthesis. In Proc. 41st International Conference on Machine Learning Vol. 235 (PMLR, 2024).

63. Dhariwal, P. & Nichol, A. Diffusion Models Beat GANs on Image Synthesis. In Advances in Neural Information Processing Systems 34 8780–8794 (Curran Associates, 2021).

64. Otsu, N. A Threshold Selection Method from Gray-Level Histograms. IEEE Trans. Syst. Man Cybern. 9, 62–66 (1979).

65. Kuhn, H. W. The Hungarian method for the assignment problem. Nav. Res. Logist. Q. 2, 83–97 (1955).

66. Everingham, M., Van Gool, L., Williams, C. K. I., Winn, J. & Zisserman, A. The pascal visual object classes (VOC) challenge. Int. J. Comput. Vis. 88, 303–338 (2010).

67. Kliper Gross, O., Rotem, O. & Zaritsky, A. DM4ISL: high-fidelity and uncertainty-aware in silico labeling of fluorescence images via diffusion models. BioImage Archive 10.6019/S-BIAD3820 (2026).

68. Hartley, M. et al. The BioImage Archive - building a home for life-sciences microscopy data. J. Mol. Biol. 434, 167505 (2022).

69. Rotem, O., Kliper Gross, O. & Zaritsky, A. DM4ISL: trained models, train/test splits, and example data. Preprint at 10.5281/ZENODO.21921959 (2026).

